# GSTT1 mediates stemness and FGFR inhibitor sensitivity in pancreatic cancer through regulation of CD133 (*PROM1*)

**DOI:** 10.1101/2025.08.06.668944

**Authors:** Deborah de la Caridad Delgado Herrera, Alejandro Arroyo Roman, Riyan N. Campbell, Emily Lasse-Opsahl, Carrie McCracken, Lisa Sadzewicz, Luke Tallon, Elana J. Fertig, Christina M. Ferrer

## Abstract

Pancreatic ductal adenocarcinoma (PDA) is among the deadliest malignancies, driven by metastatic progression and profound cellular heterogeneity. We previously identified glutathione S-transferase theta 1 (GSTT1) as a regulator of a slow-cycling, highly metastatic tumor cell population, suggesting that GSTT1^High^ cells may possess stem-like properties. Here, we define the functional and molecular features of this subpopulation in metastatic PDA. Using a mCherry-tagged *Gstt1* reporter system in metastatic murine PDAC cells, we enriched for Gstt1^High^ cells and observed increased tumor sphere formation, accompanied by upregulation of stemness-associated genes including *PROM1* (CD133) and activation of Wnt and FGF signaling pathways. In human PDA models, CD133^High^GSTT1^High^ cells exhibited enhanced tumor sphere initiation and expansion compared to other populations, defining a maximal stem-like state. Notably, sensitivity to FGFR inhibitors was observed only under tumor sphere conditions, highlighting a context-dependent therapeutic vulnerability. Mechanistically, FGFR3 expression correlated with GSTT1 and CD133 levels, and FGF signaling was required to sustain this state. *GSTT1* knockdown reduced CD133 protein levels, impaired tumor sphere formation, and altered sensitivity to FGFR inhibition. These findings were largely recapitulated in patient-derived PDA organoids, where *GSTT1* and *PROM1* co-expression predicted increased tumor sphere formation and enhanced response to the multi-kinase inhibitor Nintedanib. Together, these results identify a GSTT1^High^CD133^High^ stem-like subpopulation in metastatic PDA and identify an FGFR-dependent signaling axis that sustains this state, representing a potential therapeutic vulnerability.

## INTRODUCTION

Pancreatic ductal adenocarcinoma (PDA) remains one of the most lethal malignancies, with a 5-year survival rate of approximately 8% in patients with metastatic disease (refs. 22, 46). Most patients present with advanced tumors, and although combination chemotherapies such as FOLFIRINOX provide modest benefit (refs. 6, 16), metastatic progression remains the principal cause of mortality. A deeper understanding of the cellular heterogeneity within metastatic lesions is therefore critical.

Stem-like tumor cell populations have been implicated in tumor initiation, metastasis, and therapeutic resistance in PDA. However, stemness in pancreatic cancer is not defined by a single marker, and the identity and regulation of stem-like cells within established metastatic tumors remain incompletely understood. Markers including CD24, CD44, ALDH activity, and CD133 (PROM1) have been associated with tumor-initiating capacity in subsets of pancreatic cancers (refs. 2,3,15,23,27). Among these, CD133 has been linked to tumorigenicity and poor clinical outcomes (refs. 15, 28, 29), yet how CD133-positive cells are maintained within metastatic lesions remains unclear.

We previously identified glutathione S-transferase theta 1 (GSTT1) as uniquely required for metastatic dissemination in PDA while dispensable for primary tumor growth (ref. 10). Within established metastatic lesions, GSTT1^High^ tumor cells represent a slow-cycling, EMT-enriched subpopulation with heightened metastatic capacity (refs. 10). Given the established association between EMT, reduced proliferation, and stem-like states (refs. 45, 53), these findings raise the possibility that GSTT1 may mark or regulate a stemness-associated program within metastatic tumors. However, whether GSTT1 functionally contributes to stem-like behavior, and how such a state is maintained, remains unknown.

Here, we demonstrate that GSTT1^High^ metastatic tumor cells are enriched for a stemness-associated transcriptional program, including *PROM1* (CD133). Functionally, we show that CD133 and GSTT1 cooperatively define a hierarchical stem-like state, with CD133^High^GSTT1^High^ cells exhibiting the most robust tumor sphere–forming capacity. Mechanistically, we identify FGFR3–STAT3 signaling as a regulator of this program and demonstrate that disruption of this pathway impairs tumor-sphere growth. Finally, we show that this stem-like state is associated with selective sensitivity to FGFR inhibition, revealing a context-dependent therapeutic vulnerability. Together, these findings define a distinct stem-like subpopulation within metastatic PDA and identify an FGFR-dependent signaling axis that sustains this state.

## RESULTS

### Gstt1^High^ Metastatic Cells are Enriched with Stem Cell Characteristics

Our previous work identified the glutathione S-transferase, *GSTT1,* as a novel mediator of metastasis. Importantly, within metastatic lesions, Gstt1^high^ cells represent a highly metastatic, slow-cycling population with EMT features. Additionally, we demonstrated that *Gstt1* is required for metastatic tumor sphere growth *in vitro* (ref. 10). Tumor sphere culture conditions serve as a selective environment that enhances the enrichment of cancer stem and progenitor cells (ref. 8, 21). To further understand metastatic heterogeneity and determine whether the Gstt1^high^ metastatic subpopulation has higher stemness properties, we utilized our previously generated Cas9 system where the endogenous mouse *Gstt1* locus was tagged to a *mCherry* reporter (ref. 10) (**Fig. 1A, left panel**). Lung metastatic-derived PDAC cell lines from the *p48-Cre/p53^F/+^Kras^G12D/+^*(*KPC*) mouse model were previously isolated (ref. 10) and generated to express this construct. Cell populations were sorted based on mCherry expression as previously described (ref. 10), and mCherry^High^ and mCherry^Low^ cell populations were then cultured under serum-free, nonadherent tumor sphere conditions for 10 days to enrich for the cancer stem cell population. After 10 days in culture, tumor spheres were analyzed for cell growth and subjected to bulk RNA-Seq analysis for identification of differential gene expression pathways (**Fig. 1A, right panel**). Both tumor sphere size and number can be used to characterize cancer stem/progenitor cell populations within *in vitro* cultured cancer cells (ref. 21). Quantification of tumor spheres at day 10 revealed an increase in tumor sphere number (**Fig. 1B, 1C**) and size (**Fig. 1B, 1D**) in mCherry^High^ sorted populations, indicating Gstt1 promotes enhanced self-renewal capacity. Western blot and flow cytometry analysis demonstrated that tumor spheres retained differential Gstt1 (**Fig. 1E**) and mCherry (**Fig. S1A, S1B**) expression after 10 days in culture. To identify unique gene expression signatures in these Gstt1^high^ metastatic spheres, Day 10 tumor sphere populations were then subjected to bulk RNA-Seq. Unbiased gene expression profiling identified n=106 significantly differentially upregulated and n=12 downregulated genes (>2FC) in mCherry^High^ compared to mCherry^Low^ tumor spheres. Interestingly, mCherry^High^ tumor spheres demonstrated enrichment in various markers commonly associated with pancreatic cancer stemness and aggressiveness (*Aldh1a1*, *Adh1*, *Lgr5*, and *Prom1*) (ref. 15, 33) as well as Wnt signaling (*Wnt4*, *Klf12*) (ref. 43) (**Fig. 1F**). Pathway analysis of differentially enriched gene signatures using DAVID (**Fig. 1G**) and IPA (**Fig. S1C**) demonstrated mCherry^High^ upregulated gene signatures consistent with glutathione S-transferase activity (oxidoreductase activity, xenobiotic metabolism, Phase I functionalization of compounds) and downregulation of pathways involved in cell differentiation, consistent with a more stem-like phenotype. Together, these results underscore the role of Gstt1 in metastatic stemness and identify a gene signature that may promote the self-renewal phenotype observed in Gstt1^high^ metastatic cells.

**Figure 1.**
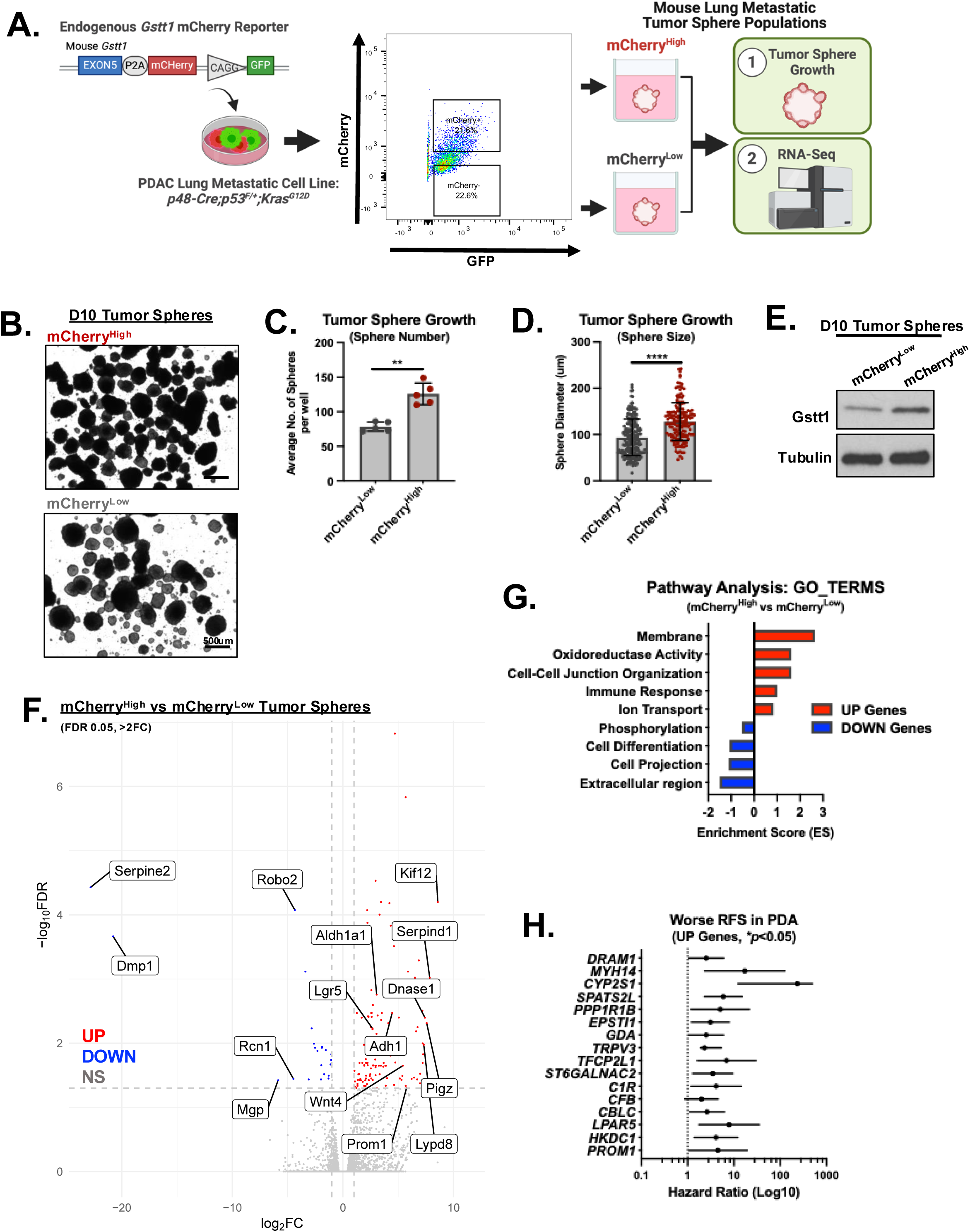
Gstt1^High^ Metastatic Tumor Spheres are Enriched with Stem Cell Signatures Associated with Poor Relapse-Free Survival in PDA. (A) PDAC-derived lung metastatic cells were generated to express *GFP+* and *mCherry* from the endogenous *Gstt1* mouse locus. Top (20%) mCherry^High^ and bottom (20%) mCherry^Low^ tumor cell populations were sorted using fluorescence-activated cell sorting (FACS) and grown as tumor spheres for 10 days. Tumor spheres were then subjected to bulk RNA-Seq. Created with Biorender.com. (B) Representative brightfield images of tumor sphere growth in mCherry populations. 500µm scale. (C) Quantification of the average number of tumor spheres per well. Data represent N=5 independent experiments, error bars indicate mean ± s.d.. Two-sided *t-test* with Welch’s correction was used to determine statistical significance between groups. (D) Quantification of tumor sphere diameter. Each data point represents a single sphere from n=5 independent experiments. Data are represented as mean s.d. Two-sided *t-test* with Welch’s correction was used to determine statistical significance between groups. (E) Western blot demonstrating Gstt1 levels in Day 10 tumor sphere populations from (B-D). (F) Volcano plot depicting differentially expressed genes (DEGs) in mCherry^High^ compared to mCherry^Low^ tumor spheres (FDR 0.05, >2FC). Stemness-associated genes are annotated in red (UP genes) and in blue (DN genes). (G) DAVID pathway analysis 106 upregulated genes and 12 downregulated genes (GO_TERMs, unpaired, two-sided *t-test* using Bonferroni correction). (H) Forest plot depicting hazard ratio (HR, Log10) of upregulated tumor sphere signature genes demonstrating poor relapse-free survival (RFS) in pancreatic cancer patients (PDA) using KMplot. Data represents relapse-free survival (RFS) in N=177 patient dataset based on the mean expression of selected genes. A Mantel-Haenszel log-rank test was used to determine significance, *\*P*<0.05.

### Identifying Gstt1-Associated Stemness Signatures in Human Pancreatic Cancer

We next aimed to investigate the relevance of the mouse Gstt1^High^ tumor sphere signature to human metastatic pancreatic cancer. Utilizing the human Cancer Cell Line Encyclopedia (CCLE) database (https://depmap.org/), we applied the Gstt1^High^ upregulated gene signature (n=106) to all pancreatic cancer cell lines (**Fig. S2A, left panel**). We first filtered out genes that were either absent in human pancreatic cancer cell lines or showed low expression, concentrating primarily on the expression of these genes in metastatic-derived (liver) and ascites-derived pancreatic cancer cell lines (**Fig. S2A, middle panel**). Next, to assess the relevance of this newly identified stemness gene signature in human pancreatic cancer, we investigated the effect of the expression of individual genes on overall survival outcomes in pancreatic cancer patients (N=63 genes) (**Fig. S2A, right panel**). From this analysis, we identified 37 genes that were predictive of survival outcomes in patients with pancreatic disease – 21 genes where high expression was associated with poor overall survival and 16 genes where high expression was associated with better overall survival (**Fig. S2B**). For the 21 genes where high expression was associated with poor overall survival, 16 of these were also significantly associated with poor relapse-free survival (**Fig. 1H**), indicating that this stemness-associated gene signature may play a role in tumor recurrence in pancreatic cancer patients. From these results, we focused on candidate genes involved in cancer stemness, i.e., *PROM1,* and pinpointed human cell line models that could be used as tools to interrogate the functional role of these genes in GSTT1-mediated stemness and heterogeneity in metastatic cells.

### GSTT1^High^ Metastatic Pancreatic Cancer Cell Lines Exclusively Express *PROM1*/CD133 and Contain High Tumor Sphere Forming Potential

*PROM1* encodes the pentosan membrane glycoprotein, CD133, and is one of the most well-characterized biomarkers used for the identification and isolation of cancer stem cells (CSCs) (ref. 32). CD133 has been shown by various groups to be a functional marker of tumor-initiating cells in pancreatic cancer (ref. 2, 7, 15) and has been linked to invasiveness and metastatic potential (ref. 7, 29). Consistent with these reports, we find that high *PROM1* expression is associated with poor overall survival in pancreatic cancer (**Fig. 2A, top panel**). Interestingly, we also find a striking association between high *PROM1* expression and poor relapse-free survival in pancreatic cancer patients (**Fig. 2A, bottom panel**), suggesting a larger role for *PROM1* in tumor recurrence and chemotherapy resistance, a key feature of cancer stem cells.

**Figure 2.**
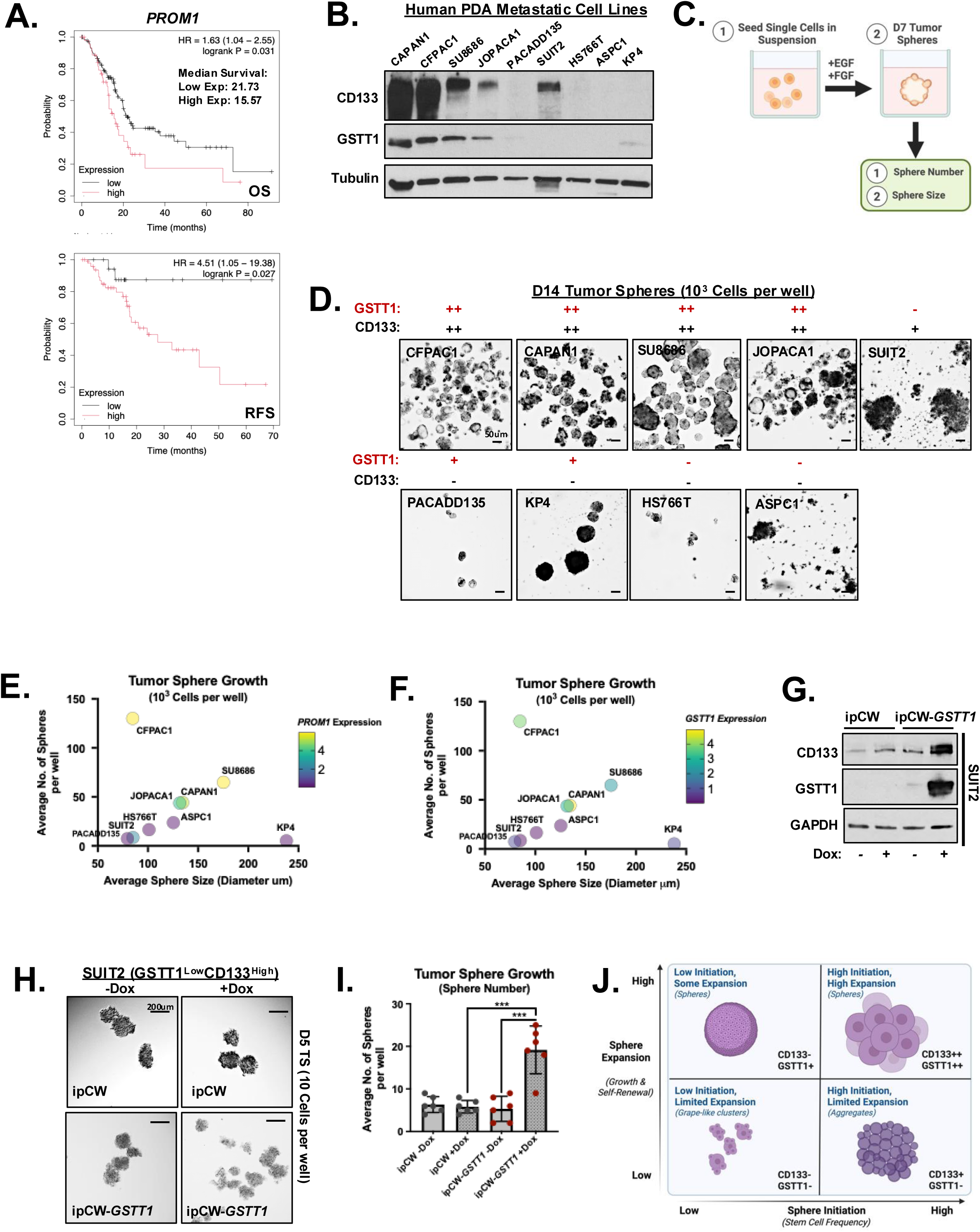
*PROM1* is Co-expressed with *GSTT1* in a Subset of Metastatic Human Pancreatic Cancer Cell Lines and is Associated with High Tumor Sphere Formation Capacity. (A) Kaplan–Meier analysis of *PROM1* expression in pancreatic cancer patients showing overall survival (OS, top) and relapse-free survival (RFS, bottom) using KMplot (N = 177 patients, stratified by mean gene expression). Significance was determined using a Mantel–Haenszel log-rank test (\**P* < 0.05). (B) Western blot analysis of CD133 and GSTT1 protein levels in whole-cell lysates from a panel of human metastatic pancreatic cancer cell lines, N=9. Data are representative of at least three independent experiments. (C) Schematic of tumor sphere culture conditions for the pancreatic cancer cell lines shown in (B). Created with BioRender.com. (D) Representative bright-field images of tumor spheres at day 14 from cell lines plated at 1 × 10³ cells per well and stratified by CD133 and GSTT1 expression. Images are representative of N=3 independent experiments. Scale bar, 50 μm. (E) Quantification of average tumor sphere number per well (y-axis) and average tumor sphere size (x-axis) at day 14 across N=9 cell lines. Cell lines are color-coded based on *PROM1* expression. (F) Same analysis as in (E), with cell lines color-coded based on *GSTT1* expression. *PROM1* and *GSTT1* mRNA expression values represent data from Supplemental Fig. 3A–B. Data represent N = 3 independent experiments with three technical replicates each. (G) SUIT2 cells expressing inducible ipCW control or ipCW-*GSTT1* were treated with doxycycline for 48 hours and analyzed by western blot using the indicated antibodies. (H) Representative bright-field images of tumor spheres derived from SUIT2 conditions in (G) at day 5 post-seeding. Scale bar, 200 μm. (I) Quantification of tumor sphere growth from (H). Data represent N = 3 independent experiments with three technical replicates each, error bars indicate mean ± s.e.m. (J) Schematic model summarizing findings from tumor sphere initiation and expansion assays, illustrating the hierarchical roles of CD133 and GSTT1. Created with BioRender.com.

Tumor recurrence due to chemotherapy resistance is largely attributed to the presence of tumor-initiating CSCs, and in pancreatic cancer, these treatments have been shown to enrich the population of CD133+ CSCs (ref. 15). Interestingly, CD133+ CSCs are more resistant to chemotherapy and radiation, which can lead to tumor recurrence (ref. 39). Glutathione S-transferases (GSTs), such as GSTT1, also function in the development of resistance to standard antineoplastic therapies through glutathionylation and subsequent detoxification of chemotherapeutic compounds (ref. 34, 47, 50). To examine the relationship between *PROM1*/CD133 and GSTT1 in metastatic PDAC, we analyzed a panel of human pancreatic cancer cell lines derived from metastatic sites (https://depmap.org/). Quantitative PCR and immunoblotting identified four cell lines that co-expressed CD133 and GSTT1 (CFPAC1, SU8686, Capan1, JOPACA1), four lines that lacked expression of both markers (ASPC1, PacaDD135, HS766T, KP4), and one CD133⁺GSTT1⁻ line (SUIT2) (**Fig. 2B, Fig. S3A-S3C**).

We next assessed whether these expression patterns correlated with functional stemness using tumor sphere assays, which enrich for stem-like cells based on clonal growth under non-adherent, serum-free conditions (ref. 21, 26, 56) (**Fig. 2C**). When plated at bulk density (1 × 10³ cells), CD133^High^GSTT1^High^ cell lines formed significantly more spheres than CD133^Low^GSTT1^Low^ lines (**Fig. 2D-2F**). Moreover, CD133^High^GSTT1^High^ cells predominantly formed round, compact spheroids, whereas CD133^Low^GSTT1^Low^ cells formed irregular, grape-like aggregates (**Fig. 2D**), a morphology previously associated with reduced therapeutic resistance (ref. 37). CD133^High^GSTT1^High^ lines also trended toward larger average sphere size (**Fig. 2E, 2F**), indicating enhanced post-initiation growth capacity.

To thoroughly investigate initiation versus post-initiation properties, we performed limiting dilution assays in all cell lines at 1, 10, and 100 cells per well. At single-cell dilution, CD133^High^ cell lines exhibited the highest frequency of positive wells, consistent with enrichment for bona fide sphere-initiating cells (**Fig. S3D, S3E**). At 10 cells per well, differences in initiation and expansion become more pronounced. CD133^High^GSTT1^High^ lines displayed the highest proportion of positive wells (≥75–85%) and formed numerous, well-organized spheres (**Fig. S3D, S3F**). Notably, KP4 cells, which lack CD133 but express low levels of GSTT1, generated fewer positive wells yet consistently formed very large spheres, suggesting strong proliferative capacity despite lower initiation capacity (**Fig. S3D, S3F**). Notably, SUIT2 cells (CD133^High^GSTT1^Low^) retained the ability to initiate growth but formed only small, poorly organized aggregates, indicating that CD133 expression is sufficient for initiation but insufficient for organized sphere formation (**Fig. S3D, S3F**). At 100 cells per well, initiation was no longer limiting and most cell lines formed spheres in nearly all wells, with marked differences in sphere number, size, and morphology. CD133^High^GSTT1^High^ cells formed the most numerous and structurally organized spheres, KP4 cells retained large sphere size, and SUIT2 cells continued to form aggregates rather than true spheres (**Fig. S3D, S3G**). Consistent with these functional differences, flow cytometric analysis of stemness-associated surface markers revealed that CD133^Low^GSTT1^Low^ cell lines were enriched for CD44⁺ cells and exhibited reduced CD24 expression, whereas CD133⁺ cell lines displayed a substantially higher proportion of CD24⁺ cells, suggesting that these cells are more likely to co-express additional markers associated with stem-like behavior (**Fig. S3H-I**).

To directly test whether GSTT1 promotes sphere expansion and organization, we overexpressed *GSTT1* in CD133^High^GSTT1^Low^ SUIT2 cells (**Fig. 2G**). While SUIT2 cells initiated growth at single-cell dilution (**Fig. S3D, S3E**), *GSTT1* overexpression significantly increased sphere number and promoted a more organized, sphere-like morphology at 10 cells per well (**Fig. 2H, 2I**). Notably, *GSTT1* overexpression also increased CD133 expression in SUIT2 cells (**Fig. 2G**), suggesting that GSTT1 reinforces a CD133-associated stem-like state.

Collectively, these findings demonstrate that GSTT1^High^ human cell lines recapitulate the *Prom1* enrichment observed in mouse Gstt1^High^ tumor spheres (**Fig. 1B**) and support a hierarchical model in which CD133 governs sphere initiation, whereas GSTT1 drives post-initiation expansion, sphere number, and structural organization, with the most robust stem-like phenotype confined to CD133⁺GSTT1⁺ populations (**Fig. 2J**).

### CD133^High^GSTT1^High^ Metastatic Tumor Spheres are Sensitive to FGFR Inhibitors

As previously noted, CD133 is a well-established marker of self-renewing CSCs in various solid tumors, and CD133⁺ cells are widely recognized for their resistance to standard chemotherapy and radiotherapy (ref. 32). Compounds such as metformin (ref. 30) and curcumin (ref. 20) have been shown to reduce CD133 expression in hepatocellular carcinoma. However, aside from CD133-targeted CAR-T therapies in glioblastoma (ref. 54), limited efforts have been made to specifically target CD133⁺ tumor cells, particularly in pancreatic cancer.

To identify compounds capable of selectively targeting CD133^High^ cells, we utilized the Genomics of Drug Sensitivity in Cancer (GDSC) database within the DepMap portal (https://depmap.org/). Metastatic pancreatic cancer cell lines were stratified based on *PROM1* (CD133) expression, and compound sensitivity from the Sanger GDSC1 screen was analyzed in relation to *PROM1* levels. This screen identified BIBF-1120 (Nintedanib) as the top compound exhibiting increased sensitivity in *PROM1^High^*pancreatic cancer cell lines (**Fig. 3A**). BIBF-1120 is a multi-kinase inhibitor that targets VEGFR, PDGFR, and FGFR (ref. 17). Although primarily studied for idiopathic pulmonary fibrosis (IPF), it has demonstrated preclinical efficacy in reducing primary tumor growth in pancreatic cancer models (ref. 24). However, its potential activity against CSCs, particularly CD133⁺ cells, remains unexplored.

**Figure 3.**
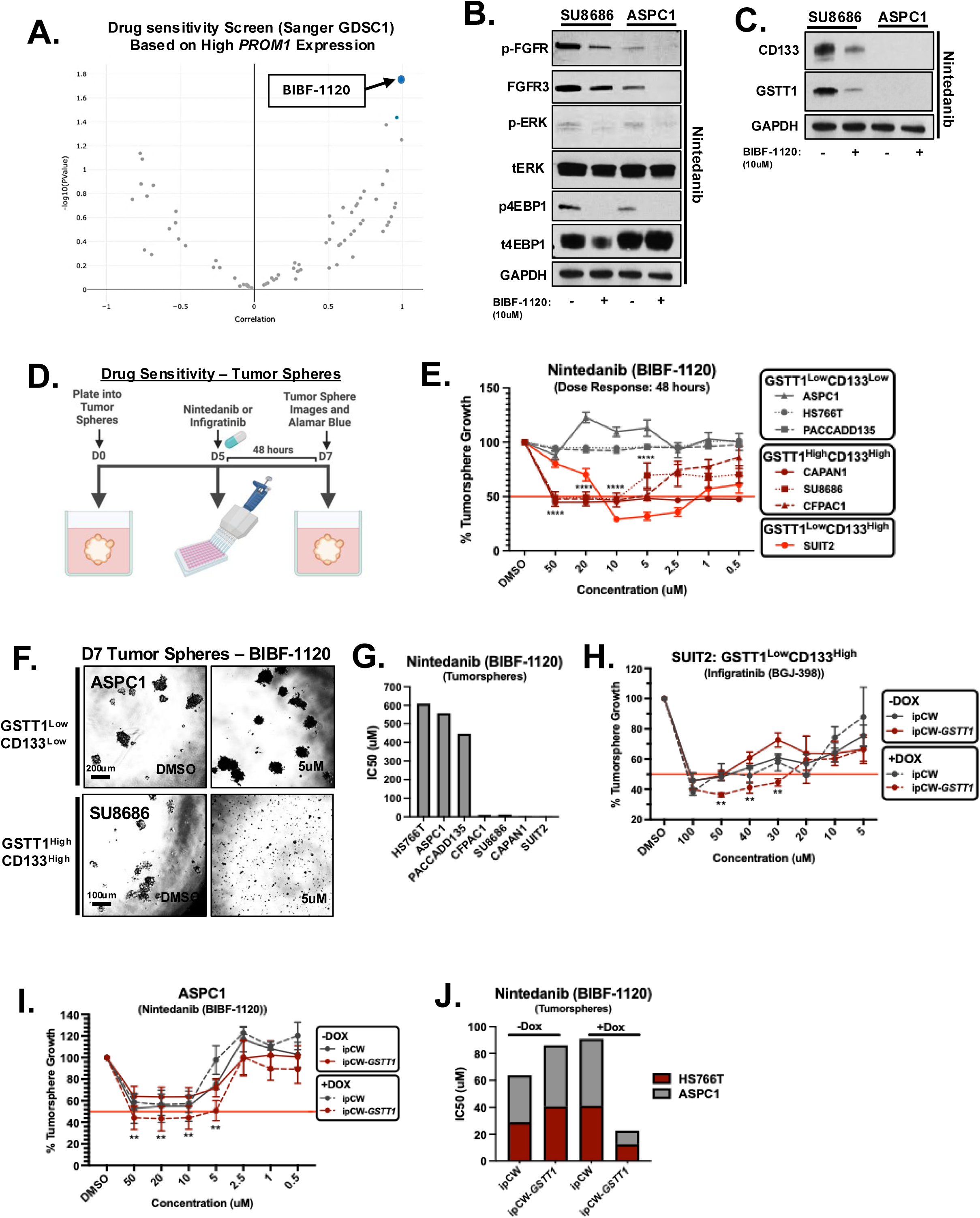
GSTT1 potentiates Nintedanib sensitivity in CD133⁺ metastatic pancreatic tumor spheres. (A) Unbiased analysis of the Sanger GDSC1 drug sensitivity screen (CCLE), stratified by *PROM1* expression, identified BIBF-1120 as a candidate compound selectively sensitizing *PROM1*^High^ cells. Drug–gene associations are shown as −log10 *P*-values, with a correlation coefficient approaching 1. (B,C) GSTT1^Low^CD133^Low^ and GSTT1^High^CD133^High^ metastatic pancreatic cancer cell lines were treated with DMSO or 10 µM BIBF-1120 (Nintedanib). Whole-cell lysates were subjected to western blot analysis using the indicated antibodies. The same lysates were used in panels B and C. (D) Schematic illustrating tumor sphere drug response assays. Human metastatic pancreatic cancer cell lines were plated under low-attachment conditions in 96-well format and allowed to form tumor spheres for 5 days prior to treatment with BIBF-1120 (Nintedanib) or BGJ-398 (Infigratinib). After 48 hours of treatment, Alamar Blue was added to assess tumor sphere viability. Created with BioRender.com. (E) Tumor sphere growth in response to escalating doses of BIBF-1120, stratified by GSTT1 and CD133 expression status. Data are presented as percent of DMSO control for each cell line. Data represent N = 3 independent experiments with three technical replicates each, error bars indicate standard deviation (s.d.). Statistical significance between GSTT1^High^CD133^High^ and GSTT1^Low^CD133^Low^ cell lines was assessed at each dose using a two-sided t-test with Welch’s correction (\*\*\*\**P* < 0.0001). (F) Bright-field (BF) representative images of Day 7 ASPC1 and SU8686 tumor spheres treated with DMSO or 5 µM BIBF-1120. Scale bars, 100–200 µm. (G) Calculated IC₅₀ values for BIBF-1120 (µM) across metastatic tumor sphere cell lines. IC₅₀ values were derived from the average of N = 3 independent experiments. (H) SUIT2 tumor spheres were grown for 5 days, induced with doxycycline to express ipCW control or ipCW-*GSTT1* and then treated with escalating doses of BIBF-1120. Results are shown as percentage of DMSO control. Data represent N = 3 independent experiments with three technical replicates each, error bars indicate mean ± s.e.m. Statistical significance was determined using a two-sided t-test with Welch’s correction, comparing the ipCW-*GSTT1* + Dox condition to all other conditions at each dose (\*\**P* < 0.01). (I) Day 5 ASPC1 tumor spheres induced with doxycycline to express ipCW control or ipCW-*GSTT1* were treated with escalating doses of BIBF-1120. Results are shown as percent of DMSO control. Data represent N = 3 independent experiments with three technical replicates each, error bars indicate mean ± s.e.m. Statistical significance was assessed as in (H) (\*\**P* < 0.01). (J) IC₅₀ values for each condition in ASPC1 and HS766T tumor spheres, calculated from the average of N = 3 independent experiments.

To investigate this, we first performed dose-response assays using BIBF-1120 across eight pancreatic cancer cell lines (**Fig. S4A**). Although Western blot analysis confirmed a reduction in downstream RTK signaling (both phosphorylated FGFR and ERK) (**Fig. 3B**) and notably, a reduction in both GSTT1 and CD133 expression (**Fig. 3C**) we observed no differential antiproliferative effects between CD133^High^GSTT1^High^ and CD133^Low^GSTT1^Low^ cell lines under standard 2D cell culture conditions (**Fig. S4A**). One challenge in CSC-targeted drug discovery is that conventional cell culture conditions fail to recapitulate the growth environment of CSCs (ref. 26). Tumor sphere cultures, however, provide a more representative *in vitro* platform for screening CSC-targeted therapies (ref. 26). Given this, we cultured pancreatic cancer cell lines in low-attachment plates with tumor sphere media for five days, followed by treatment with increasing concentrations of BIBF-1120. After an additional five days, cell viability was assessed using the resazurin-based Alamar Blue assay (**Fig. 3D**). Under tumor sphere conditions, CD133^High^ cells exhibited marked sensitivity to BIBF-1120, while CD133^Low^ cells were largely unresponsive (**Fig. 3E**, **Fig. 3F**, **Fig. 3G**). These results emphasize the importance of tumor sphere models when screening CSC-targeting therapies and suggest that BIBF-1120 may selectively target signaling pathways essential for the survival and proliferation of CD133^High^GSTT1^High^ metastatic tumor cells.

As noted, BIBF-1120 targets multiple receptor tyrosine kinases (RTKs), including the FGFR family. FGFRs are highly conserved transmembrane kinases that activate pathways such as MAPK and AKT, which are frequently dysregulated in cancer (ref. 1). FGF/FGFR signaling is critical for sphere-forming capacity in pancreatic cancer cell lines (ref. 40), and our previous work identified *FGFR3* as essential for tumor sphere growth in metastatic-derived pancreatic cancer cells (ref. 10). To explore whether CD133^High^GSTT1^High^ tumor spheres are differentially sensitive to FGFR inhibitors, CD133^High^GSTT1^High^ SU8686 cells and CD133^Low^GSTT1^Low^ KP4 cells were treated with the selective small molecule pan-FGFR inhibitor BGJ-398 (Infigratinib). BGJ-398 treatment led to reduced phospho-FGFR and, interestingly, total FGFR3 levels, along with suppression of downstream effectors, including phospho-ERK and phospho-4EBP1 (**Fig. S4B**). Notably, FGFR inhibition also decreased CD133 and GSTT1 protein levels in SU8686 cells (**Fig. S4C**), recapitulating our results in BIBF-1120 treated cells (**Fig. 3C**). Similar to BIBF-1120, no differential response was observed under 2D cell culture conditions (**Fig. S4D**), but under tumor sphere conditions, two CD133^High^GSTT1^High^ cell lines (SU8686 and Capan1) showed dose-dependent sensitivity to BGJ-398, whereas the three CD133^Low^GSTT1^Low^ cell lines (ASPC1, HS766T, PACADD135) showed no response (**Fig. S4E**). Interestingly, SUIT2 cells, which express high CD133 but low GSTT1, responded only at higher BGJ-398 concentrations, suggesting that GSTT1 may contribute to FGFR inhibitor sensitivity (**Fig. S4E**). Consistent with this, *GSTT1* overexpression enhanced BGJ-398 sensitivity in SUIT2 cells, indicating that both GSTT1 and CD133 are required for a complete response to FGFR inhibitors (**Fig. 3H**). Together, these findings support CD133 as a biomarker for identifying GSTT1⁺ metastatic pancreatic cancer cells that may be responsive to FGFR inhibitors.

Given this selective sensitivity, we next asked whether ectopic expression of *GSTT1* could confer drug sensitivity in CD133^Low^GSTT1^Low^ cell lines. Using a doxycycline-inducible system, we stably transduced ASPC1 and HS766T cells with either an empty vector or *GSTT1*. Three days of doxycycline induction led to *GSTT1* overexpression (**Fig. S4F, S4G**). Dose-response assays revealed that *GSTT1* overexpression rendered CD133^Low^GSTT1^Low^ tumor spheres sensitive to both BIBF-1120 and BGJ-398 in a dose-dependent manner (**Fig. 3I**, **Fig. 3J; Fig. S4H–J**). These results indicate that *GSTT1* overexpression is sufficient to sensitize tumor spheres to FGFR-targeting therapies.

### FGF–FGFR3–STAT3 Signaling Regulates the GSTT1-CD133 Stem-Like Program in Metastatic PDA Cells

Tumor sphere culture provides a widely used system for enriching cancer stem–like cell (CSC) populations from heterogeneous tumor samples, based on the ability of individual stem-like cells to survive and proliferate under non-adherent, serum-free conditions supplemented with defined growth factors such as EGF and FGF that support self-renewal pathways (ref. 21, 23, 26) (**Fig. 4A**). Consistent with enrichment of stem-like populations under these conditions, CD133^High^GSTT1^High^ metastatic pancreatic cancer cell lines (CFPAC1 and SU8686) cultured under tumor sphere conditions exhibited a marked increase in both GSTT1 and CD133 protein expression compared to standard 2D adherent culture (**Fig. 4B**). These observations suggested that signals present in tumor sphere media promote activation of the CD133–GSTT1 program.

**Figure 4.**
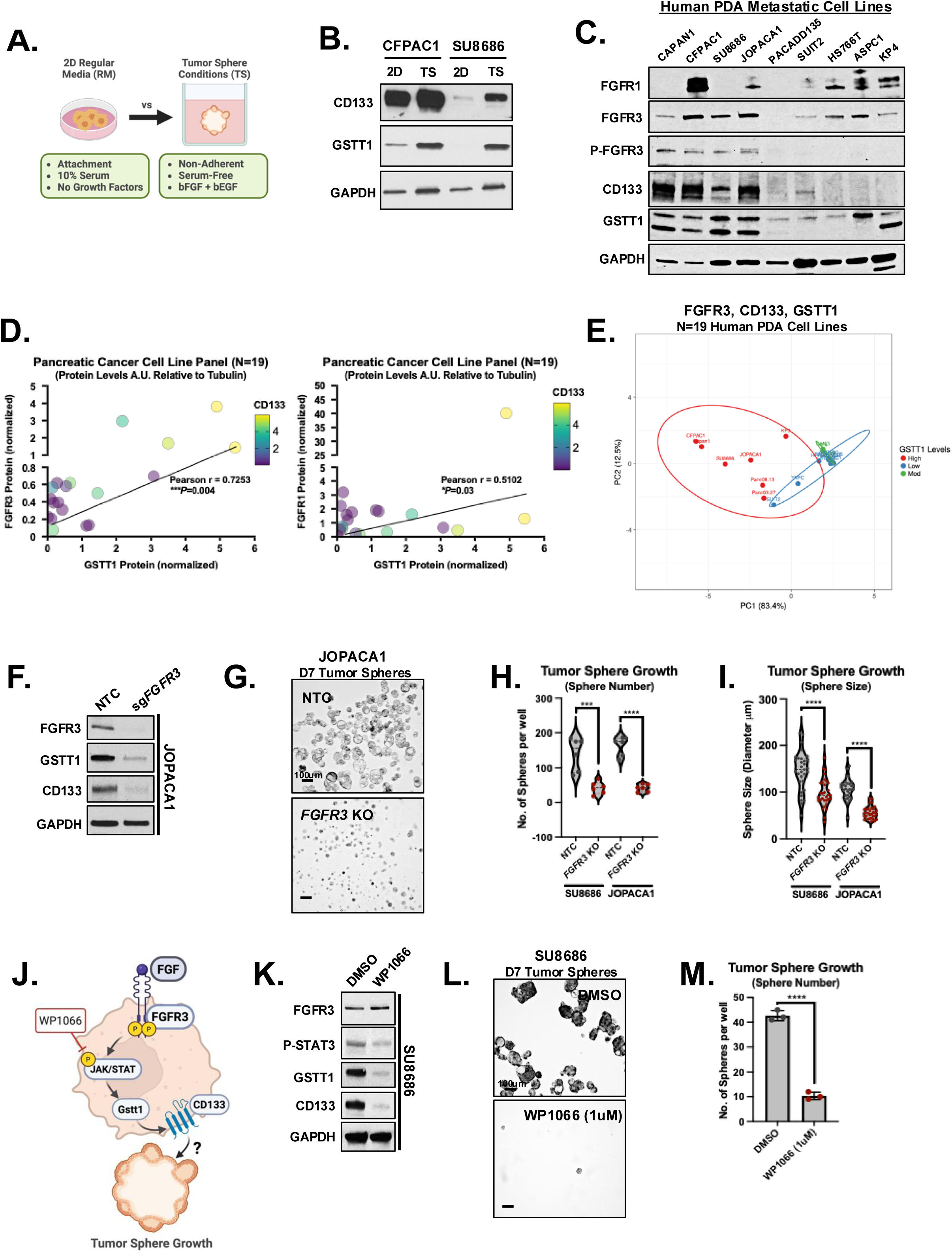
FGF-FGFR Signaling Drives CD133 and GSTT1 Enrichment in Tumor Sphere Conditions. (A) Schematic depicting differences between 2D attachment culture and low-attachment tumor sphere conditions. Created with BioRender. (B) Western blot showing CD133 and GSTT1 protein levels in whole-cell lysates from CFPAC1 and SU8686 metastatic cell lines grown under 2D attachment or low-attachment tumor sphere conditions for 7 days. (C) Western blot of CD133, GSTT1, FGFR1, FGFR3, and phospho-FGFR3 (Y724) in a panel of human metastatic pancreatic cancer cell lines (N=9). (D) Western blot quantification of GSTT1 versus FGFR3 (left) and GSTT1 versus FGFR1 (right) across N = 19 pancreatic cancer cell lines. Heatmap coloring indicates CD133 protein levels. All protein values are normalized to tubulin or GAPDH loading controls. Quantification represents N = 3 independent experiments. (E) Principal component analysis (PCA) of N = 19 pancreatic cancer cell lines using normalized protein expression of GSTT1, CD133, and FGFR3, showing clustering patterns based on these markers. Color coding indicates GSTT1 levels. Created with Clustvis. (F) Western blot of FGFR3, GSTT1, CD133, and GAPDH in JOPACA1 cells expressing either non-targeting control (NTC) or CRISPR-mediated *FGFR3* knockout (KO) using three pooled guides. (G) Brightfield images of tumor sphere growth in NTC or *FGFR3* KO JOPACA1 cells 10 days post-plating. (H, I) Quantification of the number of tumor spheres per well (H) and tumor sphere size (diameter, µm) (I) for each condition and cell line. Each data point represents a single sphere across N=3 independent experiments in JOPACA1 and SU8686 NTC or *FGFR3* KO cells. Data represent N = 3 replicates from N = 3 independent experiments per cell line. Statistical significance determined by two-sided t-test with Welch’s correction (\*\*\**P* < 0.001, \*\*\*\*\**P* < 0.0001). (J) Schematic depicting the JAK/STAT inhibitor WP1066 and the experimental hypothesis. Created with BioRender.com. (K) Western blot of indicated proteins in SU8686 cells treated with DMSO or 1 μM WP1066 for 48 hours. (L) Brightfield images of tumor sphere growth in SU8686 cells treated with DMSO or 1 μM WP1066 10 days post-plating. (M) Quantification of the number of tumor spheres per well in SU8686 cells. Data represent N = 3 replicates from N = 3 independent experiments, error bars indicate standard deviation (s.d). Statistical significance determined by two-sided t-test with Welch’s correction (\*\*\*\*\**P* < 0.0001).

To determine whether soluble factors within tumor sphere media were sufficient to induce this phenotype independent of 3D architecture, we cultured CD133^High^GSTT1^High^ SU8686 cells under adherent conditions using either standard media or tumor sphere media, with or without FGFR inhibitors (**Fig. S5A**). Tumor sphere media alone significantly increased CD133 surface expression even under adherent conditions, reaching levels comparable to those observed in 3D tumor spheres (**Fig. S5B–C**). These culture conditions also increased GSTT1 and CD133 protein expression together with phosphorylation of FGFR3 downstream effectors STAT3 and ERK (**Fig. S5D**). Importantly, pharmacologic FGFR inhibition suppressed these effects, demonstrating that factors present in tumor sphere media are sufficient to induce GSTT1 and CD133 expression through FGF-FGFR-mediated signaling.

Because fibroblast growth factors signal through fibroblast growth factor receptors (FGFRs) to regulate pathways involved in proliferation, survival, and stem cell maintenance, we next examined which FGFR family members might account for this effect. FGFR3 protein expression was specifically associated with GSTT1 and CD133 expression across metastatic pancreatic cancer cell lines (**Fig. 4C**). In contrast, FGFR1 showed no correlation with CD133 or GSTT1 expression, and FGFR2 expression was minimal in this panel (**Fig. 4C** and data not shown). Notably, phosphorylated FGFR3 was detected exclusively in CD133^High^GSTT1^High^ cells, indicating active FGFR3 signaling specifically within this stem-like subpopulation (**Fig. 4C**). Consistent with this observation, SUIT2 cells, which express low levels of CD133 but lack GSTT1, did not exhibit detectable phospho-FGFR3, suggesting that FGFR3 activation rather than expression alone distinguishes the tumor sphere–competent CSC-like subset. To further evaluate this association, we expanded the analysis to a larger panel of human pancreatic cancer cell lines (N = 19). FGFR3 protein levels significantly correlated with GSTT1 expression (Pearson r = 0.7253, p = 0.004) and showed a similar association with CD133 (**Fig. 4D-4E** and **Fig. S5E**). In contrast, FGFR1 showed a weaker correlation with GSTT1 expression (Pearson r = 0.5102, p = 0.03), while FGFR2 expression was minimal (**Fig. 4D** and **Fig. S5E**), supporting FGFR3 as the primary FGFR associated with the GSTT1-positive stem-like state.

To directly test whether FGFR3 is required for maintenance of this phenotype, we generated *FGFR3* knockout cells in CD133^High^GSTT1^High^ metastatic pancreatic cancer cell lines JOPACA1 (**Fig. 4F**) and SU8686 (**Fig. S5G**). Loss of *FGFR3* significantly impaired tumor sphere growth (**Fig. 4G–I** and **Fig. S5F**) and was accompanied by decreased expression of both CD133 and GSTT1 (**Fig. 4F** and **Fig. S5G**), confirming that FGFR3 signaling is necessary to sustain the CSC-like state. These findings are consistent with our previous results where *Fgfr3* was also required for tumor sphere growth in in metastatic *KPC* cell lines (ref.10), further supporting a conserved role for FGFR3 in maintaining stem-like properties in metastatic pancreatic cancer.

We next investigated the downstream signaling pathways responsible for this effect. Pharmacologic interrogation of canonical FGFR effector pathways revealed that inhibition of STAT3 with WP1066 markedly reduced CD133 and GSTT1 expression and suppressed tumor sphere formation, phenocopying both FGFR inhibition and *FGFR3* genetic ablation (**Fig. 4J–M** and **Fig. S5H–J**). In contrast, inhibition of MEK or ERK failed to suppress CD133 or GSTT1 expression and instead resulted in a compensatory increase in STAT3 phosphorylation accompanied by increased CD133 expression (data not shown). Together, these findings define an FGFR3–STAT3 signaling axis that maintains the CD133–GSTT1 stem-like program in metastatic pancreatic cancer cells.

Finally, to determine whether GSTT1 alone is sufficient to drive this tumor sphere phenotype, we overexpressed *GSTT1* in CD133^Low^GSTT1^Low^ metastatic pancreatic cancer cell lines (ASPC1 and HS766T) under both FGF-supplemented and FGF-depleted conditions using a doxycycline-inducible system (**Fig. S6A, S6B**). *GSTT1* overexpression modestly increased CD133 protein levels, but only in the presence of FGF, and this partial upregulation did not translate into significant tumor sphere formation (**Fig. S6A–C**). CD133 levels in *GSTT1*-overexpressing cells remained substantially lower than those observed in GSTT1^High^CD133^High^ CFPAC1 cells (**Fig. S6E**). Notably, *GSTT1* overexpression in these CD133^Low^ cells did induce FGFR3 phosphorylation and STAT3 activation (**Fig. S6D–E**), providing a mechanistic explanation for their enhanced sensitivity to FGFR inhibitors despite limited functional stem-like growth. In contrast, in SUIT2 cells, which possess basal CD133 expression but lack GSTT1, overexpression of *GSTT1* robustly increased p-FGFR3, elevated CD133, and enhanced tumor sphere formation (**Fig. S6F, Fig. 2G–I**). Together, these results indicate that GSTT1 alone is insufficient to induce tumor sphere formation in the absence of basal CD133 expression, but can potentiate stem-like growth in cells with pre-existing CD133. Collectively, these findings support a model in which FGF–FGFR3 signaling establishes a stem-like signaling framework, while GSTT1 amplifies this program to sustain CD133 expression, tumor sphere expansion, and FGFR inhibitor sensitivity.

### *GSTT1* is Required to Maintain CD133 Expression, Tumor Sphere Formation and Sensitivity to FGFR Inhibitors

Next, we sought to determine whether GSTT1 was required to maintain CD133 levels and subsequent tumor sphere formation. To test this question, we knocked down *GSTT1* using shRNA in three GSTT1^High^CD133^High^ metastatic cell lines (SU8686, CFPAC1 and Capan1). qPCR and western blotting demonstrate a significant reduction in *GSTT1* mRNA (SU8686) and protein expression in shRNA-expressing cells compared to cells expressing a non-targeting control (**Fig. S7A, Fig. 5A**). However, *GSTT1* knockdown resulted in decreased CD133 protein levels without significant changes in *PROM1* mRNA expression, indicating post-transcriptional regulation (**Fig. S7A)**. Immunoprecipitation experiments revealed no detectable glutathione modification of CD133 in response to glutathione addition in the form of N-acetylcysteine (NAC) (**Fig. S7B**), whereas treatment with the proteasome inhibitor MG132 rescued CD133 protein levels in *GSTT1*-depleted cells, supporting a role for GSTT1 in regulating CD133 protein stability rather than transcription or direct glutathione modification (**Fig. S7C)**. Interestingly, although *GSTT1* overexpression increased FGFR3 protein levels and phosphorylation in the presence of FGF in CD133^Low^GSTT1^Low^ cells (**Fig. S6D-S6E**), FGFR3 remained unchanged upon *GSTT1* knockdown (**Fig. 5A** and **Fig. S7C**), further supporting our model that FGFR3–STAT3 acts upstream to initiate the stem-like program, with GSTT1 functioning downstream to amplify and stabilize CD133-dependent tumor sphere formation.

**Figure 5.**
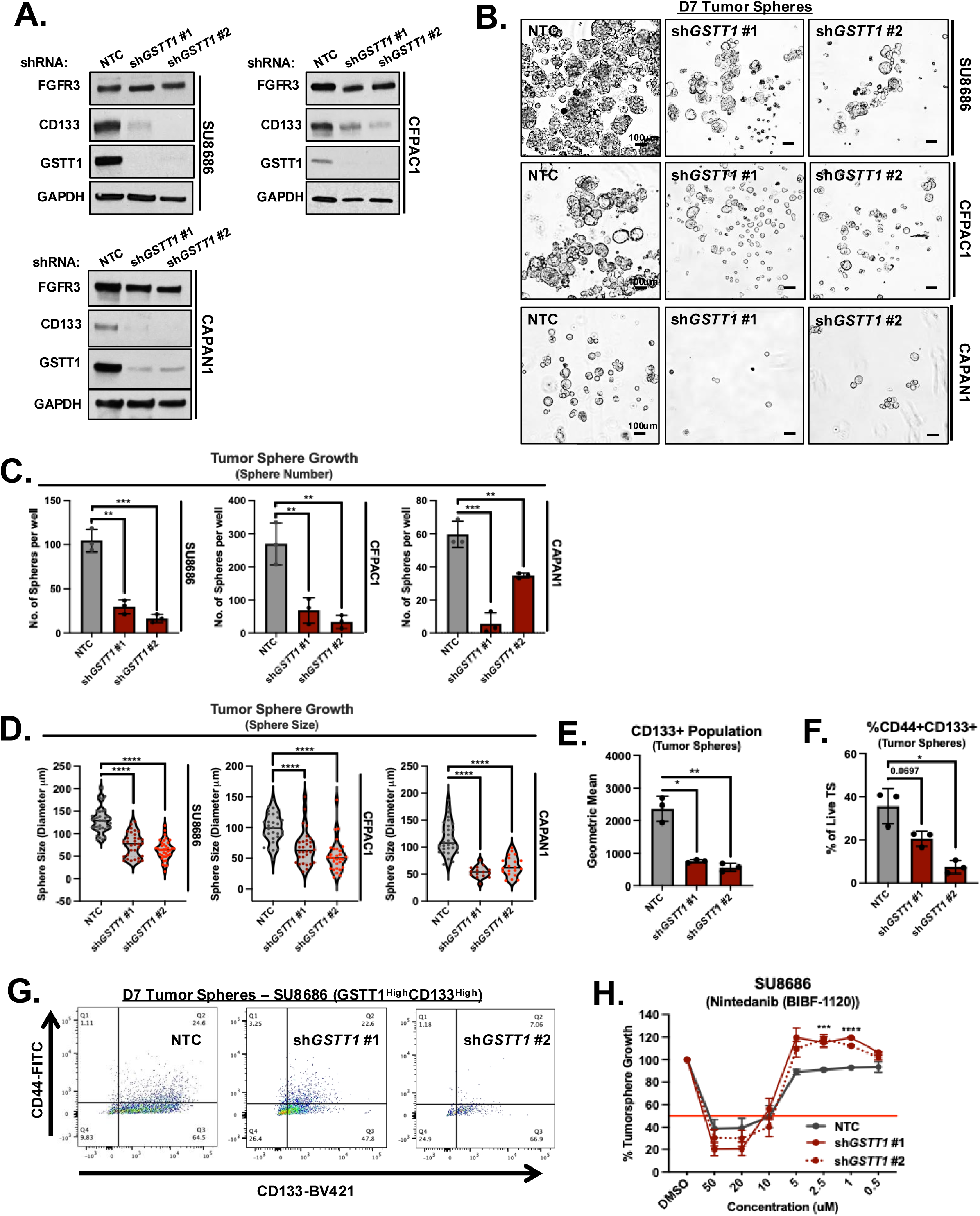
GSTT1 is Required to Maintain CD133 Expression, Tumor Sphere Formation and Sensitivity to FGFR Inhibitors. (A) Whole-cell lysates from SU8686, CFPAC1 and CAPAN1 (GSTT1^High^CD133^High^) cells expressing either NT Control vector or two independent *GSTT1* shRNAs were subjected to western blotting using indicated antibodies. (B) Bright-field (BF) representative images of Day 7 tumor spheres from each condition. Scale bar, 100µm. (C) Bar plot depicting the quantification of the number of tumor spheres per well for each condition. Data represent N=3 replicates from N=3 independent experiments, error bars indicate standard deviation (s.d). Two-sided *t-test* with Welch’s correction was used to determine statistical significance between groups (\*\**P*<0.01, \*\*\**P*<0.005). (D) Violin plot depicting tumor sphere size (diameter, µm) for each condition and cell line. Each data point represents a single sphere across N=3 independent experiments. Two-sided *t-test* with Welch’s correction was used to determine statistical significance between groups, (\*\**P*<0.01, \*\*\*\**P*<0.0001). (E) Flow cytometry analysis of CD133+ cells based on surface marker expression in SU8686 tumor spheres expressing either NT Control vector or two independent *GSTT1* shRNAs. Data are presented as geometric mean (GeoMean), error bars indicate standard deviation (s.d). Two-sided *t-test* with Welch’s correction was used to determine statistical significance between groups (\**P*<0.05, \*\**P*<0.01). (F,G) Flow cytometry analysis of CD44+CD133+ cell population based on surface marker expression in SU8686 tumor spheres expressing either NT Control vector or two independent *GSTT1* shRNAs. Data represented as CD144+CD133+ population relative to the live tumor sphere population. Data represent N=3 replicates from N=3 independent experiments, error bars indicate standard deviation (s.d). Two-sided *t-test* with Welch’s correction was used to determine statistical significance between groups (\**P*<0.05). (H) SU8686 (GSTT1^High^CD133^High^) cells expressing either NT Control vector or two individual *GSTT1* shRNAs were cultured as tumor spheres for 5 days and subsequently subjected to a dose response of BIBF-1120 (Nintedanib) for 48 hours. Results are shown as % tumor sphere growth relative to each DMSO treated control for each condition. Data represent N=3 replicates from N=3 independent experiments, error bars indicate standard deviation (s.d). Two-sided *t-test* with Welch’s correction was used to determine statistical significance between NT Control and each shRNA at both 2.5 μM and 1 μM dose (\*\**P*<0.01, \*\*\**P*<0.005).

Next, we tested whether this decrease in CD133 levels was associated with a reduction in tumor sphere formation. Consistent with reduced CD133 protein (**Fig. 5A**), *GSTT1* knockdown decreased both tumor sphere number and size across all three cell lines (**Fig. 5B–D**), indicating that GSTT1-mediated regulation of CD133 protein is required to maintain tumor sphere phenotypes.

To complement tumor sphere formation, we next assessed the surface expression of CD133 and the widely used stemness marker CD44 in tumor spheres expressing either control shRNA or *GSTT1*-targeting shRNAs. Similar to our results above, knockdown of *GSTT1* in SU8686 cells resulted in a reduction in CD133 surface marker expression and CD133⁺ tumor spheres across both shRNA clones (**Fig. 5E**). However, only one *GSTT1*-targeting shRNA significantly decreased both the CD44⁺CD133⁺ double-positive tumor sphere population (**Fig. 5F**, **Fig. 5G**). These results suggest that CD133 may act in concert with other stemness markers to maintain tumor sphere formation in GSTT1+ pancreatic cells.

Our results indicate that CD133 expression determines sensitivity to FGFR inhibitors in GSTT1⁺ cells. To further investigate this, we tested whether a reduction in CD133 protein following *GSTT1* knockdown would affect sensitivity to FGFR inhibitors. We performed dose-response assays in SU8686 tumor spheres, stably expressing either a non-targeting control or two independent *GSTT1* shRNAs, and treated with increasing doses of BIBF-1120 (Nintedanib). Notably, cells expressing *GSTT1*-targeting shRNAs showed reduced sensitivity to BIBF-1120 compared to the control (**Fig. 5H**). These findings indicate that GSTT1-mediated regulation of CD133 protein influences both tumor sphere formation and drug responsiveness in metastatic pancreatic cancer.

### Heterogenous *GSTT1* and *PROM1* Expression Profiles in PDA Organoids Predict Tumor Sphere Formation and Nintedanib Sensitivity

Next, we sought to determine whether patient-derived organoids exhibited a similar effect on tumor sphere formation and response to BIBF-1120 (Nintedanib) based on GSTT1 and *PROM1/*CD133 expression. To investigate this, we generated tumor spheres from pancreatic cancer organoids derived from six patients (**Fig. 6A**). RNA was isolated from these tumor spheres, and qRT-PCR was performed to assess gene expression (**Fig. 6A**, **Fig. 6B**). The tumor spheres displayed a range of *GSTT1* and *PROM1* expression levels. Notably, Organoid #2 exhibited the highest expression of both genes, while Organoids #1, #3, and #5 showed elevated *GSTT1* expression alone (**Fig. 6B**). Correlating gene expression with tumor sphere growth revealed that *GSTT1* expression was associated with increased sphere number (**Fig. 6C**, **Fig. 6D**). Furthermore, co-expression of *GSTT1* and *PROM1* correlated with increased sphere size, with Organoid #2—expressing high levels of both—producing the largest tumor spheres (**Fig. 6C** and **Fig. 6E**). These findings suggest that the observations made in human cell line models are recapitulated in patient-derived organoids, reinforcing the relevance of *GSTT1* and *PROM1* expression as potential biomarkers of a metastatic, stem-like cell population in pancreatic cancer.

**Figure 6.**
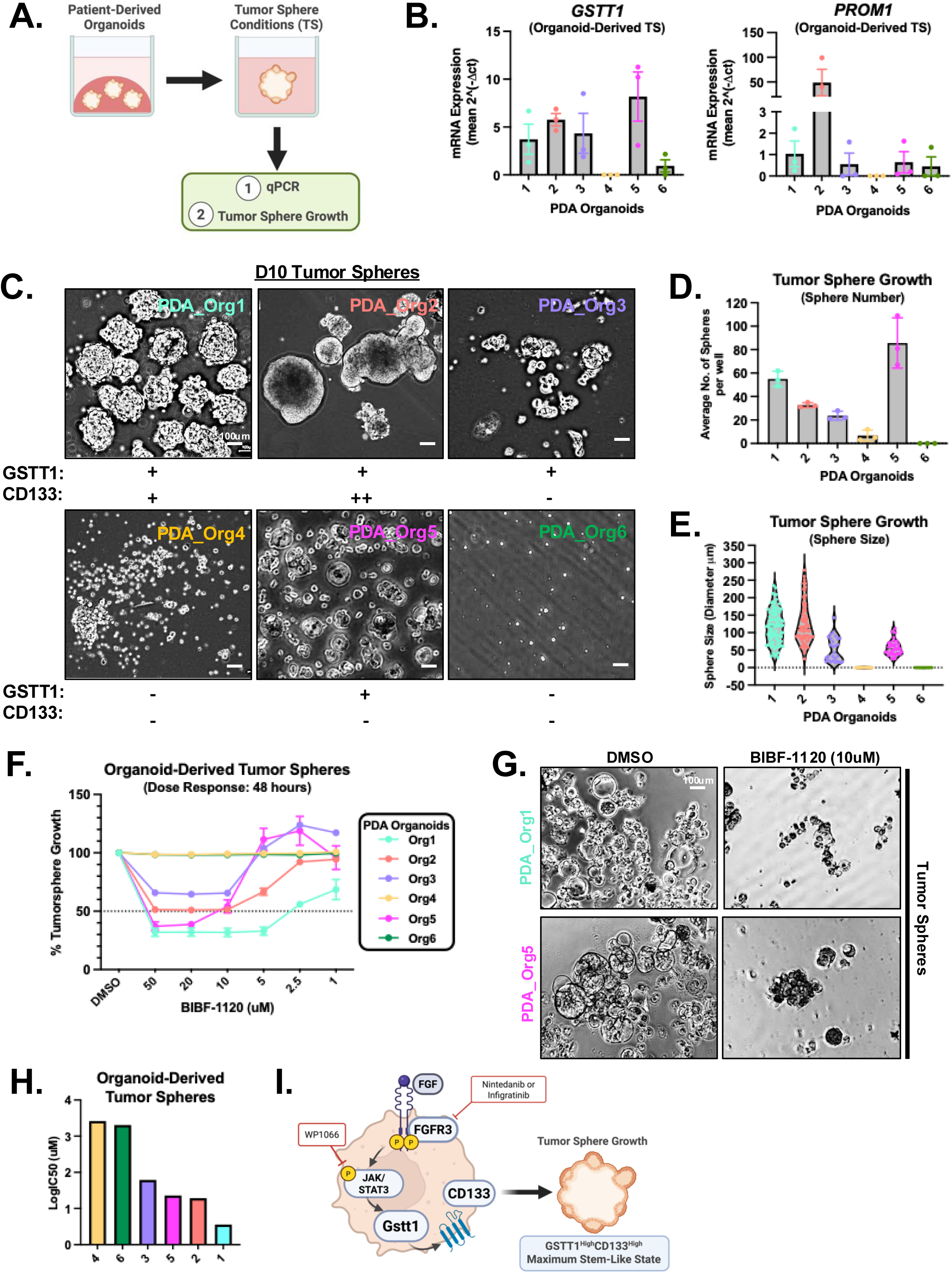
Heterogeneous Expression of *GSTT1* and *PROM1* in Patient-Derived PDA Organoids Correlates with Tumor Sphere Formation and Sensitivity to Nintedanib. (A) Schematic illustrating the workflow for culturing PDA patient-derived organoids under tumor sphere conditions, followed by qPCR analysis and tumor sphere growth assays. Created with BioRender.com. (B) qRT-PCR Expression of *GSTT1* and *PROM1* in a panel of PDA patient-derived organoid-derived tumor spheres (PDA_Org1-6). Data are represented as mean s.e.m. Data represents 3 independent experiments. (C) Representative brightfield images of tumor sphere growth in PDA organoids. Scale bar, 100 µm. (D) ImageJ quantification of the average number of tumor spheres per well. Data are represented as mean s.d. Data represent 3 independent experiments. (E) Quantification of tumor sphere size (diameter, µm). Each data point represents a single sphere across three independent experiments. (F) Dose–response of organoid-derived tumor spheres to BIBF-1120 (Nintedanib). Results are shown as percentage of DMSO control for each organoid line. Data represent N=2 independent experiments, with n=3 replicates each, error bars indicate standard deviation (s.d). (G) Representative bright-field images showing tumor sphere growth in PDA organoids treated with 10 µM BIBF-1120. Scale bar: 100 µm. (H) LogIC₅₀ values for organoid-derived tumor spheres treated with BIBF-1120. (I) Schematic model depicting overall findings. Created with BioRender.

Next, we investigated whether expression of *GSTT1* and *PROM1* in organoid-derived tumor spheres could predict sensitivity to the pan-FGFR inhibitor BIBF-1120 (Nintedanib). To assess this, we performed dose-response experiments by culturing pancreatic organoids under tumor sphere-forming conditions. Organoids were dissociated into single cells and seeded into 96-well low-attachment plates containing tumor sphere media. After 5 days of sphere formation, cultures were treated with varying concentrations of BIBF-1120 and allowed to grow for an additional 5 days. Cell viability was measured on Day 10 using the resazurin-based Alamar Blue assay. Interestingly, we observed that tumor spheres derived from Organoids #1, #2, and #5 exhibited the greatest sensitivity to FGFR inhibition by Nintedanib (**Fig. 6F**, **Fig. 6G**). In contrast, and consistent with our findings in human PDA cell lines, Organoids #4 and #6 were resistant to BIBF-1120, consistent with a high IC50 value (**Fig. 6H**) that correlated with their limited tumor sphere-forming ability and low expression of *GSTT1* and *PROM1*. Collectively, these findings indicate that patient-derived GSTT1^High^ tumor spheres are selectively sensitive to FGFR inhibition by BIBF-1120, suggesting that GSTT1 may serve as a predictive biomarker for targeting aggressive, stem-like populations in pancreatic cancer with FGFR inhibitors. Collectively, this model positions GSTT1 as both a biomarker and functional mediator of metastatic pancreatic CSC plasticity, and identifies the FGFR3–STAT3–GSTT1–CD133 axis as a therapeutically targetable vulnerability in metastatic PDA (**Fig. 6I**).

## DISCUSSION

Pancreatic ductal adenocarcinoma (PDA) remains highly lethal due to metastatic progression, therapeutic resistance, and extensive intratumoral heterogeneity (ref. 46). Within metastatic lesions, diverse tumor cell populations coexist with distinct proliferative capacities, metastatic potential, and drug sensitivities (ref. 12). Emerging evidence implicates a rare subpopulation of cancer stem cells as key drivers of metastasis, therapy evasion, and tumor relapse, adding an additional layer to this heterogeneity (ref. 15, 21).

In this study, we identify a metastatic PDA cell subpopulation defined by co-expression of GSTT1 and CD133 that exhibit enhanced stem-like behavior and tumor sphere–forming capacity. Building on our previous work showing that GSTT1 marks a slow-cycling, highly metastatic population enriched for epithelial-to-mesenchymal transition (EMT) features (ref. 10), we demonstrate that GSTT1^High^ cells also possess functional CSC properties. Using our previously developed Gstt1 reporter system (ref. 10) in metastatic murine PDAC cells, we find that both Gstt1^High^ and Gstt1^Low^ metastatic subpopulations retain stem-like capacity, but Gstt1^High^ cells are strongly enriched for stemness-associated pathways, defining a distinct, transcriptional state. By Day 10, a proportion of Gstt1^Low^ spheres regain GSTT1 expression (**Fig. 1E, S1A**), reflecting phenotypic plasticity under stem-cell enriching conditions. This dynamic reinforces the enrichment of a GSTT1-associated stem-like phenotype and its selective sensitivity to FGFR inhibition.

GSTT1 is a member of the glutathione S-transferase family and is canonically involved in catalyzing glutathione conjugation to target proteins and metabolites (ref. 34, 47, 50). Mechanistically, this enzymatic activity provides a plausible mechanism through which GSTT1 could stabilize or regulate proteins such as CD133, or modulate signaling pathways like FGFR3–STAT3. Our preliminary experiments using immunoprecipitation of endogenous CD133 in GSTT1^High^CD133^High^ cells did not detect direct glutathione modification of CD133, even in the presence of exogenous glutathione (N-acetylcysteine) (**Fig. S7B**). While lack of detection of this modification using this approach does not exclude this mechanism of regulation, it suggests that detection may require more sensitive approaches such as mass spectrometry, or that GSTT1 acts indirectly, for example, by modulating the cellular redox environment to influence protein stability or signaling.

Our functional studies demonstrate that GSTT1 alone is insufficient to induce a fully stem-like phenotype in the absence of basal CD133 expression. Instead, GSTT1 amplifies or stabilizes stem-like behavior once initiation capacity is established, consistent with a hierarchical model in which multiple cooperating pathways are required to sustain maximal CSC activity (**Fig. 2J**). Consistent with this, CD133^High^GSTT1^Low^ cells initiated growth but formed poorly organized aggregates, whereas CD133^High^GSTT1^High^ cells generate the most robust and structurally organized spheres (**Fig. S3D-G**). Together, these results support a model in which CD133 contributes to tumor sphere initiation, while GSTT1 enhances proliferative capacity and structural organization required for maximal stem-like behavior.

Mechanistically, we identify FGF–FGFR3 signaling as a key pathway sustaining this stem-like state. FGFR3 protein expression and phosphorylation strongly correlate with GSTT1 and CD133 levels, whereas mRNA expression is less predictive, suggesting post-translational regulation in metastatic PDA cells. Notably, phospho-FGFR3 is largely restricted to CD133^High^GSTT1^High^ cells, indicating that this subpopulation depends on active FGFR3 signaling. Consistent with this, *GSTT1* overexpression in CD133^Low^GSTT1^Low^ and CD133^High^GSTT1^Low^ cells increases FGFR3 phosphorylation and STAT3 activation (**Fig. S6D-F**). While *GSTT1* overexpression alone is insufficient to fully induce tumor sphere formation in the absence of high basal CD133 levels (**Fig. S6A, S6B**), the associated increase in phospho-FGFR3 provides a mechanistic explanation for the enhanced sensitivity of these cells to FGFR inhibition. Together, these findings suggest that GSTT1 promotes a signaling state characterized by elevated FGFR3 activation, creating a dependency that can be therapeutically exploited.

Our findings also underscore the strong context-dependence of this signaling network. Both GSTT1 and CD133 expression are enriched under tumor sphere culture conditions, and sensitivity to FGFR inhibition was largely restricted to these CSC-enriched environments. Tumor sphere media alone is sufficient to induce GSTT1 and CD133 expression even under adherent conditions, indicating that soluble niche factors, including FGF, can activate this program. These observations highlight the importance of modeling CSC-supportive microenvironments when evaluating therapeutic strategies targeting stem-like tumor cell populations.

Finally, our findings have potential therapeutic implications. FGFR inhibitors, including Nintedanib and Infigratinib, selectively impaired the viability of GSTT1^High^CD133^High^ pancreatic cancer cells under tumor sphere conditions. Patient-derived PDA organoids similarly exhibited heterogeneous *GSTT1* and *PROM1* expression, where co-expression predicted larger tumor spheres and enhanced sensitivity to FGFR inhibition. Together, these results define a distinct GSTT1^High^CD133^High^ metastatic cell state with enhanced stem-like properties and suggest that targeting the FGFR3–STAT3 axis as a therapeutically targetable dependency in this aggressive tumore cell population.

## MATERIALS AND METHODS

### Cell Lines

Mouse cell lines from lung and liver metastases were isolated from the *p48-Cre/p53F/+KrasL/+* (*Sirt6 WT* and *KO)* as previously described (ref. 10). Cells were cultured in RPMI 1640 supplemented with 10% fetal bovine serum and 1% penicillin (100 U/ml) streptomycin (100 Ug/ml) (Invitrogen Gibco).

Metastasis-derived human pancreatic cancer cell lines, CFPAC1 (CRL#1918), SU8686 (CRL#1837), Hs 766T (HTB#134), ASPC1 (CRL#1682), CAPAN1 (HTB#79), HPAFII (CRL-1997) were obtained from ATCC. PACADD135 and JOPACA1 were obtained from the Leibniz-Institut (DSMZ). SUIT2, KP4, Panc08.13, Panc03.27, KP3, PSN1, DAN-G, SW1990, YAPC, Colo357 and Panc1 were obtained from the Massachusetts General Hospital Cancer Center Cell Line Repository (Boston, MA). SU8686, ASPC1, SUIT2, CFPAC1, KP4, Panc08.13, Panc03.27, KP3, PSN1, DAN-G, SW1990, YAPC, Colo357 and Panc1 were cultured in RPMI 1640 supplemented with 10% fetal bovine serum and 1% penicillin (100 U/ml)/streptomycin (100 Ug/ml) (Invitrogen Gibco). CAPAN1, PACADD135, and JOPACA1 were cultured in IMDM supplemented with 20% fetal bovine serum and 1% penicillin (100 U/ml)/streptomycin (100 Ug/ml) (Invitrogen Gibco). Hs 766T were cultured in 1X DMEM supplemented with 10% fetal bovine serum and 1% penicillin (100 U/ml)/streptomycin (100 Ug/ml) (Invitrogen Gibco).

### Generation of *Gstt1-mCherry* Cell Line and Fluorescence Activated Cell Sorting

Endogenous tagging of murine *Gstt1-mCherry* PDAC lung metastatic cell lines was previously described (ref. 10). The lung metastatic-derived cell line generated to express *mCherry* from the endogenous *Gstt1* locus was cultured under 2D conditions, trypsinized, and filtered through 40µm mesh filters. DAPI (3 nm) was added to the cell suspension to negatively select live cells. mCherry^high^ and mCherry^low^ populations were gated on live populations as well as GFP expression. For all *in vitro* tumor sphere culture assays, the top 15-20% and bottom 15-20% were chosen as mCherry^high^ and mCherry^low^ populations, respectively, utilizing appropriate single-color and negative controls as previously described (Ferrer et al, 2024).

### Tumor Sphere Assay and Experimental Media Conditions

Human and Mouse metastatic-derived cell lines were plated as single-cell suspension as previously described (ref. 10, 23) in ultralow attachment plates (Corning). Cells were grown in DMEM/F12 serum-free medium supplemented with 1X B-27 (Thermo, #17504044), 20ng/ml EGF (recombinant mouse #315-09, Peprotech; recombinant human #AF-100-15, Peprotech) and 20ng/ml bFGF (recombinant mouse #450-33. Peprotech; recombinant human #100-18B, Peprotech). Single cells were plated at the indicated density for each experiment (refer to figure legend), and fresh media was added every 3 days. Tumor spheres were counted and photographed from days 6-10 using a Leica Matteo TL digital transmitted light microscope. Tumor Spheres were quantified using Image J and data represents a minimum of 3 biological replicates per group performed in triplicate for each experiment. Brightfield sphere images (2.5X, 4X magnification) were first threshold-filtered to eliminate debris and single cells. For sphere size, a minimum of 25 spheres per well were quantified, and data is represented as sphere diameter (µm) with a minimum of 3 biological replicates per group. For FGF supplementation experiments, 20ng/ml was supplemented into FGF-free tumor sphere media every other day.

For tumor sphere limiting dilution analysis, cells were plated at densities of 1, 10, 100, or 1,000 cells per well in ultra–low attachment plates (Corning) in complete tumor sphere media. For 1 and 10 cells per well, cells were plated in 96-well plates with 20–24 replicate wells per condition. For 100 and 1,000 cells per well, cells were plated in 24-well plates with 10–12 replicate wells per condition. Tumor spheres were cultured for up to 30 days. Each limiting dilution experiment was repeated a minimum of two independent biological times. Percent positive wells were calculated as the percentage of wells containing at least one tumor sphere and were plotted together with the average number of spheres per well.

### RNA Isolation from Tumor Spheres and RNA Sequencing

Cas9-*Gstt1*-*mCherry* expressing Tumor Sphere populations were enzymatically digested, serially filtered and sorted using a FACSAria II (BD Biosciences) and BD FACSDiva Software (v.8.0.2) Obtained data were analyzed by FlowJo V10. Cells were collected into RNA isolation buffer followed by RNA isolation using the RNeasy Mini Kit (Qiagen, 74104). In order to minimize ribosomal RNA background, samples were enriched for mRNA containing Poly A tails. We used a bead based enrichment kit - NEBNext® Poly(A) mRNA Magnetic Isolation Module (New England Biolabs, Ipswitch, MA). No modifications were made to the manufacture’s protocol. Strand-specific, dual unique indexed libraries for sequencing were made using the NEBNext® Ultra™ II Directional RNA Library Prep Kit for Illumina®(New England Biolabs,Ipswich, MA). Manufacturer protocol was modified by diluting adapter 1:30 and using 3 ul of this dilution. The size selection of the library was performed with SPRI-select beads (Beckman Coulter Genomics, Danvers, MA). Glycosylase digestion of adapter was included at the final amplification to avoid further cleanups. 16 cycles were used for PCR amplification. All Libraries were assessed on the GX Touch (Revity, Waltham, MA). Sequencing was performed on an Illumina NovaSeq 6000 using a 100bp paired-end run configuration.

### RNA Sequencing Analysis

RNAseq analysis was carried out by Maryland Genomics, Institute for Genome Sciences, UMSOM. Paired-end Illumina libraries were mapped to the mouse reference, Ensembl release GRCm39.110., using HiSat2 v2.1.0, using default mismatch parameters. Read counts for each annotated gene were calculated using HTSeq. The DESeq2 Bioconductor package (v1.5.24) was used to estimate dispersion, normalize read counts by library size to generate the counts per million for each gene, and determine differentially expressed genes between experimental groups. Differentially expressed transcripts with a p-value ≤ 0.05 and log_2_ fold change ≥ 1 were used for downstream analyses. Normalized read counts were used to compute the correlation between replicates for the same condition and compute the principal component analysis for all samples. Analysis of enriched functional categories among detected genes was performed using Ingenuity Pathway Analysis.

### Human Datasets

To analyze the expression of the n=106 upregulated genes found in mouse mCherry^High^ tumor spheres in human metastatic cell lines, the Cancer Cell Line Encyclopedia (CCLE, Depmap) (ref. 52) was utilized. Cell line origin (i.e., primary tumor, ascites, and metastasis) information was also obtained from CCLE, Depmap. Cancer Cell Line Encyclopedia (CCLE) was accessed on 1/2/24 from https://registry.opendata.aws/ccle. Expression of UP genes (upregulated in mCherry^High^ tumor spheres) from mouse tumor spheres were then investigated for expression levels in human cell lines. Genes demonstrating no or low expression (<1.0 log2(TPM+1)) in human cell lines were removed from subsequent analysis. N=63 genes were expressed in human pancreatic metastatic cell lines after filtering.

To determine whether the expression of the top enriched n=63 genes correlate with Overall Survival (OS) or Relapse-Free Survival (RFS) in pancreatic cancer (PDA) we utilized KMPlot (ref. 25). The dataset includes n=177 patients of all stages. Gene effect on survival is represented as “Worse” survival in patients with high overall expression and “Better” survival in patients with lower overall expression (log-rank *P* value, \**P*<0.05). N=37 genes demonstrated a differential effect on overall patient survival, and high expression of N=21 genes demonstrated worse overall survival. Of these N=21 genes, N=16 genes in human pancreatic metastatic cell lines displayed significant correlation with RFS in patients with pancreatic cancer.

### Identification of Compounds using CCLE-DepMap

To identify compounds to specifically target CD133^High^-expressing pancreatic cancer cells, we utilized CCLE, Depmap (ref. 52). Briefly, metastatic pancreatic cancer cell lines stratified by *PROM1*/CD133 expression (Expression Public 24Q2) were analyzed against the Genomics of Drug Sensitivity in Cancer (GDSC1) dataset containing data from 970 cell lines and 403 compounds. Unbiased analysis identified BIBF-1220 as highly correlated with sensitivity in cell lines with high *PROM1* expression. Current DepMap Release data, including CRISPR Screens, PRISM Drug Screens, Copy Number, Mutation, Expression, and Fusions can be found at *DepMap, Broad (2023). DepMap 23Q4 Public. Figshare+. Dataset.* https://doi.org/10.25452/figshare.plus.24667905.v2.https://depmap.org/portal.

### *In vitro* Compound Assays

For 2D attachment assays, cells were treated with escalating doses of either BIBF-1120 (Nintedanib, MedChem Express, #HY-50904), BGJ-398 (Infigratinib, SelleckChem, #S2183) or WP-1066 (SelleckChem, #S2796) for 48 hours with DMSO as a control. After 48 hours, cells were subjected to MTT assay (Sigma-Aldrich, #11465007001) and plates were read at 570nm as per manufacturer’s instructions. For tumor sphere treatment assays, cells were first grown as tumor spheres in 96-well low attachment plates (Corning), for 5 days, to allow tumor spheres to form. After 5 days, tumor spheres were treated with escalating doses of either BIBF-1120 (Nintedanib, MedChem Express, #HY-50904) or BGJ-398 (Infigratinib, SelleckChem, #S2183) for 48 hours with DMSO as a control in tumor sphere media. After 48 hours, tumor spheres were treated with Alamar Blue (Invitrogen, #DAL1025) for 24 hours and quantified as per manufacturer instructions. IC50 for each cell line was calculated using Graphpad Prism. For Immunoblotting, cells were treated with sublethal doses of each compound, as per IC50 calculations.

Stable *GSTT1* knockdown cell lines were generated as described below. Cells were treated with 5 μM MG132 or vehicle control (DMSO) for 24 hours to inhibit proteasomal degradation. Following treatment, whole-cell lysates were collected and subjected to Western blot analysis to assess protein stability and expression.

### Lentivirus Vectors and Stable Cell Lines

Lentivirus generation and infection were performed as previously described (ref. 23). Stable cell lines for shRNA knockdowns were generated by infection with the lentiviral vector pLKO.1-puro carrying shRNA sequence for scrambled (Addgene, Cambridge, MA, USA) or human *GSTT1 (NM_000853)* (sh#1 TCTGATTAAAGGTCAGCACTT; sh#2CTTTGCCAAGAAGAACGACAT). Stable cell lines were generated by puromycin selection.

To generate lentiviral vectors for doxycycline-inducible overexpression of wild-type human *GSTT1*, we replaced the CAS9 sequence in the ipCW-CAS9 vector (#50661, Addgene) by the 3X FLAG tagged CDS of TV1 of human *GSTT1* through Gibson cloning. Stable cell lines (SUIT2, ASPC1, HS766T) were generated by puromycin selection. Cell lines were treated with 1 μg ml^−1^ doxycycline for a minimum of three days to achieve overexpression.

### Generation of *FGFR3* Knockout Cells using CRISPR-Cas9

*FGFR3* knockout cell lines were generated in SU8686 and JOPACA1 pancreatic cancer cells using a CRISPR-Cas9 system. SpCas9 2NLS nuclease (300 pmol), negative control scrambled sgRNA#1, and a human *FGFR3* gene knockout kit containing three pooled sgRNAs were obtained from EditCo. Ribonucleoprotein (RNP) complexes were assembled according to the manufacturer’s instructions and nucleoporated into cells using the Lonza 4D-Nucleofector system. Following nucleoporation, cells were allowed to recover in complete growth media prior to downstream analyses. *FGFR3* knockout efficiency was validated by immunoblotting for FGFR3 protein expression before functional tumor sphere assays were performed.

### Real-Time qRT-PCR Analysis

Total RNA was isolated as described above. For cDNA synthesis, 1μg of total RNA was reverse-transcribed using the QuantiTect Reverse Transcription Kit (Qiagen). Real-time PCR was run in duplicate using 2X Universal SYBR Green Fast qPCR master mix (ABclonal, #RK21203) and real-time monitoring of PCR amplification was performed using the CFX96 Real-Time PCR (Bio-Rad). Data were expressed as relative mRNA levels or mean 2^(-dCT) normalized to the β-actin expression level in each sample. Primer sequences for human *GSTT1* (For- TTCCTTACTGGTCCTCACATCTC; Rev- TCACCGGATCATGGCCAGCA), human *PROM1*(For- ATTGGCATCTTCTATGGTTT; Rev- GCCTTGTCCTTGGTAGTGT), and human *GAPDH* (For- AATGAAGGGGTCATTGATGG; Rev- AAGGTGAAGGTCGGAGTCAA).

### Immunoblotting

Cell line and tumor sphere lysates were prepared in RIPA lysis buffer (150 mM NaCl, 1% NP40, 0.5% DOC, 50 mM Tris-HCl at pH 8, 0.1% SDS, 10% glycerol, 5 mM EDTA, 20 mM NaF and 1 mM Na3VO4) supplemented with a protease inhibitor cocktail. Whole-cell lysates were sonicated and cleared by centrifugation at 16000 x *g* for 20min at 4°C and analyzed by SDS-polyacrylamide gel electrophoresis (PAGE) and autoradiography. Proteins were analyzed by immunoblotting using the following primary antibodies: anti-Gstt1 (Abcam, ab175418; Abcam, ab199337), anti-Tubulin (Sigma-Aldrich, T6199), anti-CD133 (Abcam, ab284389), anti-phospho-ERK (Cell Signaling Technology, #4370), anti-total ERK (Santa Cruz Biotechnology, SC-93), anti-phospho-FGFR (pan, Cell Signaling Technology, #3471), anti-FGFR3 (ABclonal, #A19052), anti-phospho-FGFR3 (Abcam, ab155960), anti-phospho-Stat3 (Cell Signaling Technology, #9145), anti-phospho-4EBP1 (Cell Signaling Technology, #9451), anti-total 4EBP1 (Cell Signaling Technology, #9644). All primary antibodies were used at a 1:1000 dilution. Where indicated, western blot bands were quantified using ImageJ and represented as relative to Tubulin expression in each cell line.

### Surface Marker Expression of CD133, CD44 and CD24 via Flow Cytometry

Live 2D-attached cells and tumor spheres were dissociated into single-cell suspensions and incubated with anti-human CD133-BV421 (#566595, Biolegend), anti-human CD44-FITC antibodies (#11-0441-82, eBioscience) and anti-human CD24-APC (#311118, Biolegend) for 30 minutes at room temperature with end-over-end rotation, protected from light. Staining was performed in flow buffer consisting of 0.5% BSA and 0.05% sodium azide in 1× PBS. Following incubation, cells were centrifuged and washed three times with flow buffer. Prior to analysis, cells were stained with 7-AAD for viability assessment. Flow cytometry was performed using FACSAria II (BD Biosciences) or the Aurora CS (Cytek) and data were analyzed using FlowJo software (version 10).

### Immunoprecipitation

JOPACA1 cells were treated with or without 100 μM N-acetylcysteine (NAC) for 24 hours. Cells were lysed in RIPA buffer supplemented with protease inhibitors as described above. Lysates were sonicated and clarified by centrifugation at 16,000 × g for 20 minutes at 4°C. For immunoprecipitation, clarified lysates were incubated with magnetic Dynabeads Protein G (Invitrogen, 10007D) pre-conjugated with either anti-CD133 antibody (Abcam) or normal rabbit IgG control (Cell Signaling Technology, 2729S). Antibody–bead conjugates were incubated with lysates overnight at 4°C with constant rotation in IP buffer (RIPA containing 0.25 mM PMSF and protease inhibitor cocktail tablets; Sigma-Aldrich, 4693116001). Following incubation, beads were washed with IP buffer and immunoprecipitated proteins were resolved on 4–20% SDS-PAGE gels. Proteins were transferred and analyzed by immunoblotting using anti-CD133 or anti-GSH (D8) antibody (Santa Cruz Biotechnology).

### Patient-Derived Organoid Culture and Tumor Sphere Formation

Patient-derived pancreatic cancer organoids were obtained from the NCI Patient-Derived Models Repository (NCI-PDMR). Organoids were thawed and cultured on Matrigel domes using the OncoPro™ Tumoroid Culture Medium Kit (Thermo Fisher Scientific), supplemented with FGF-10 (Peprotech, #100-26) and Gastrin (Thermo Fisher Scientific, #30-061), following the manufacturer’s instructions. For tumor sphere generation, organoids were dissociated into single cells and seeded under low-attachment conditions in human tumor sphere medium, as described above.

### Statistical Analysis and Reproducibility

Student’s two-sided *t*-test was performed for two-group comparisons. Unequal variances were adjusted according to Welch’s correction. Multiple comparisons were performed using an ANOVA with post-hoc Holm–Sidak’s, Bonferroni or Geisser-Greenhouse adjusted *P* values. A log-rank test was used to determine significance for Kaplan-Meier analyses. Data distribution was assumed to be normal, but this was not formally tested. GraphPad Prism 8 was used for plotting graphs and statistical analysis. Statistical significance (*P* values) is displayed in Figures and Figure legends. All western blots represented have been repeated a minimum of two times.

## Data Availability Statement

All RNA–seq data that support the findings of this study will be deposited in the Gene Expression Omnibus (GEO). There is no restriction on data availability. Publicly available cell line expression data can be accessed through the Cancer Cell Line Encyclopedia DepMap Portal (https://depmap.org/portal/ccle/) Human PDA survival data were derived from KMPlot (https://kmplot.com/analysis/). All other data supporting the findings of this study are available from the corresponding author on reasonable request. In-house codes were previously used in the published works. Appropriate references to the original works are provided.

## ACKNOWLEDGEMENTS

This article was supported by funds through the National Cancer Institute - Cancer Center Support Grant (CCSG) – P30CA134274. The authors thank the Flow Cytometry Shared Service of the University of Maryland Marlene and Stewart Greenebaum Comprehensive Cancer Center for their support. This work was funded by the Maryland Department of Health’s Cigarette Restitution Fund Program (CH-649-CRF) and the National Cancer Institute (NIH-NCI) grant R00 CA252600 awarded to C.M.F.

## AUTHOR CONTRIBUTIONS

C.M.F. conceived and designed the study, performed experiments, and wrote the manuscript. D.D.H. performed experiments, contributed to study design, analyzed data, and wrote and critically revised the manuscript. A.A.R. and R.J.C. performed experiments and analyzed data. C.M. and L.S. performed experiments and analyzed data. L.T. contributed to RNA-seq study design. E.L.-O. and E.J.F. performed analysis and interpretation of human pancreatic single-cell RNA-seq datasets and contributed to the mechanistic framework of the study.

---

**Supp Fig 1.**
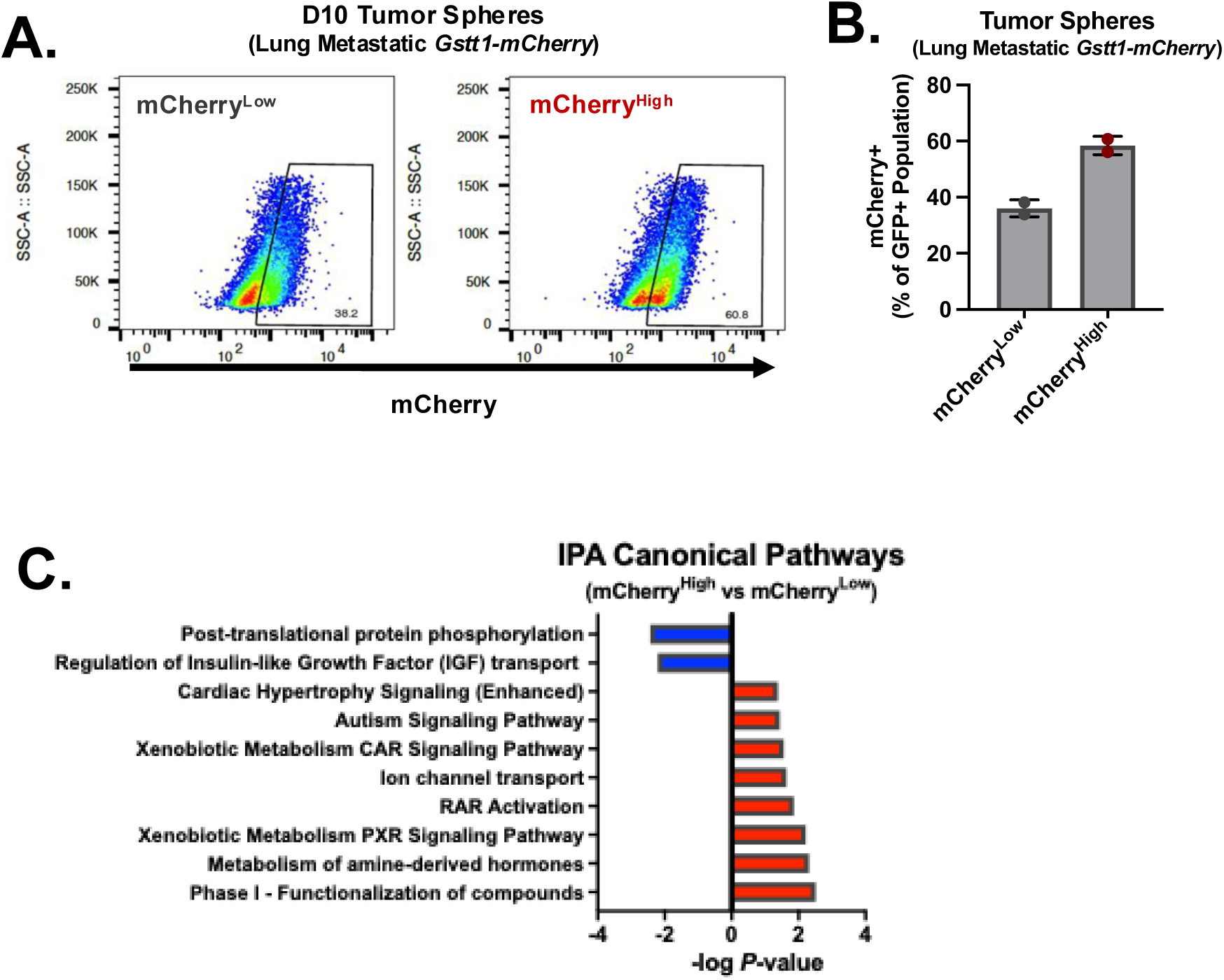

**Supp Fig 2.**
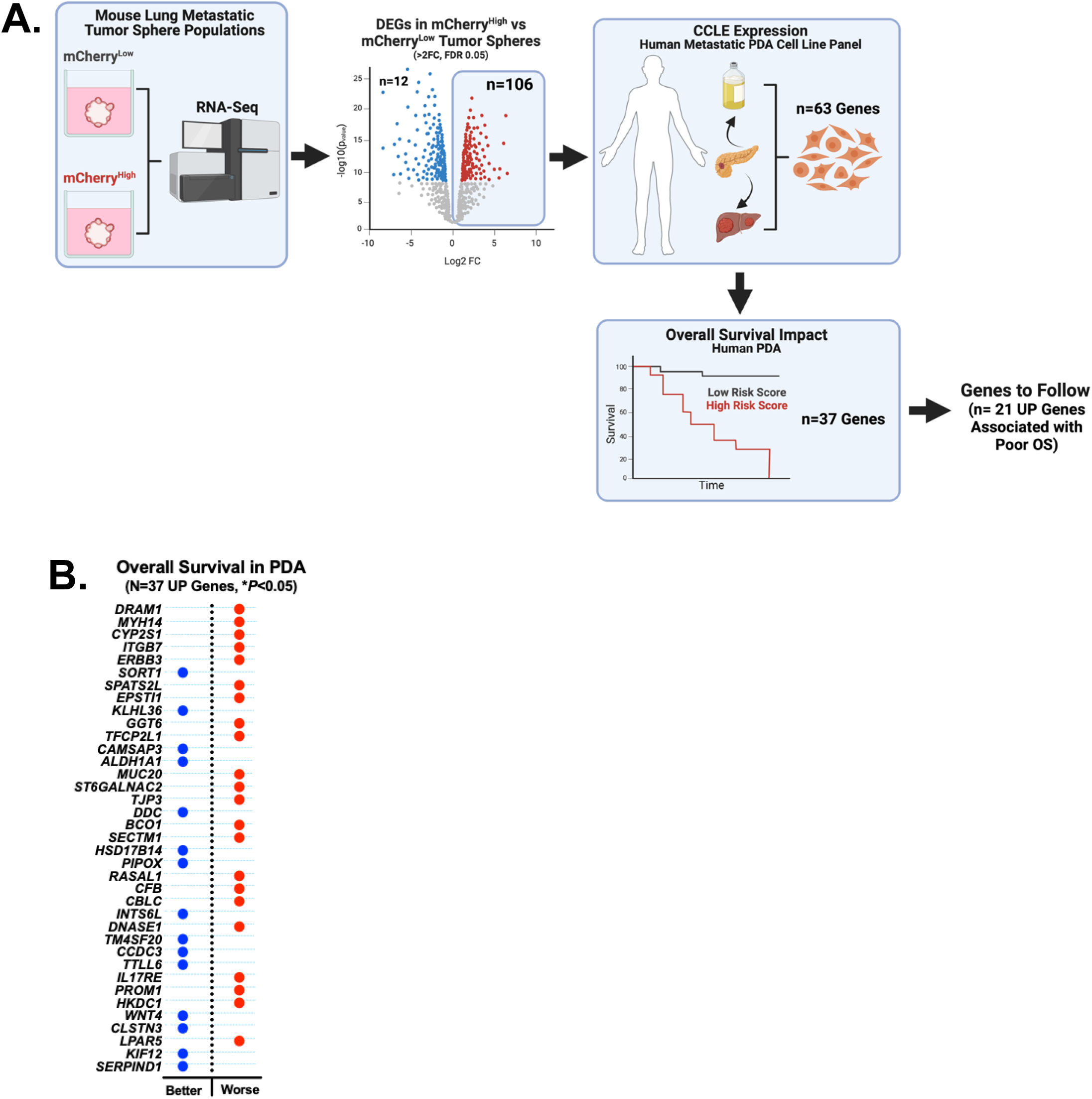

**Supp Fig 3.**
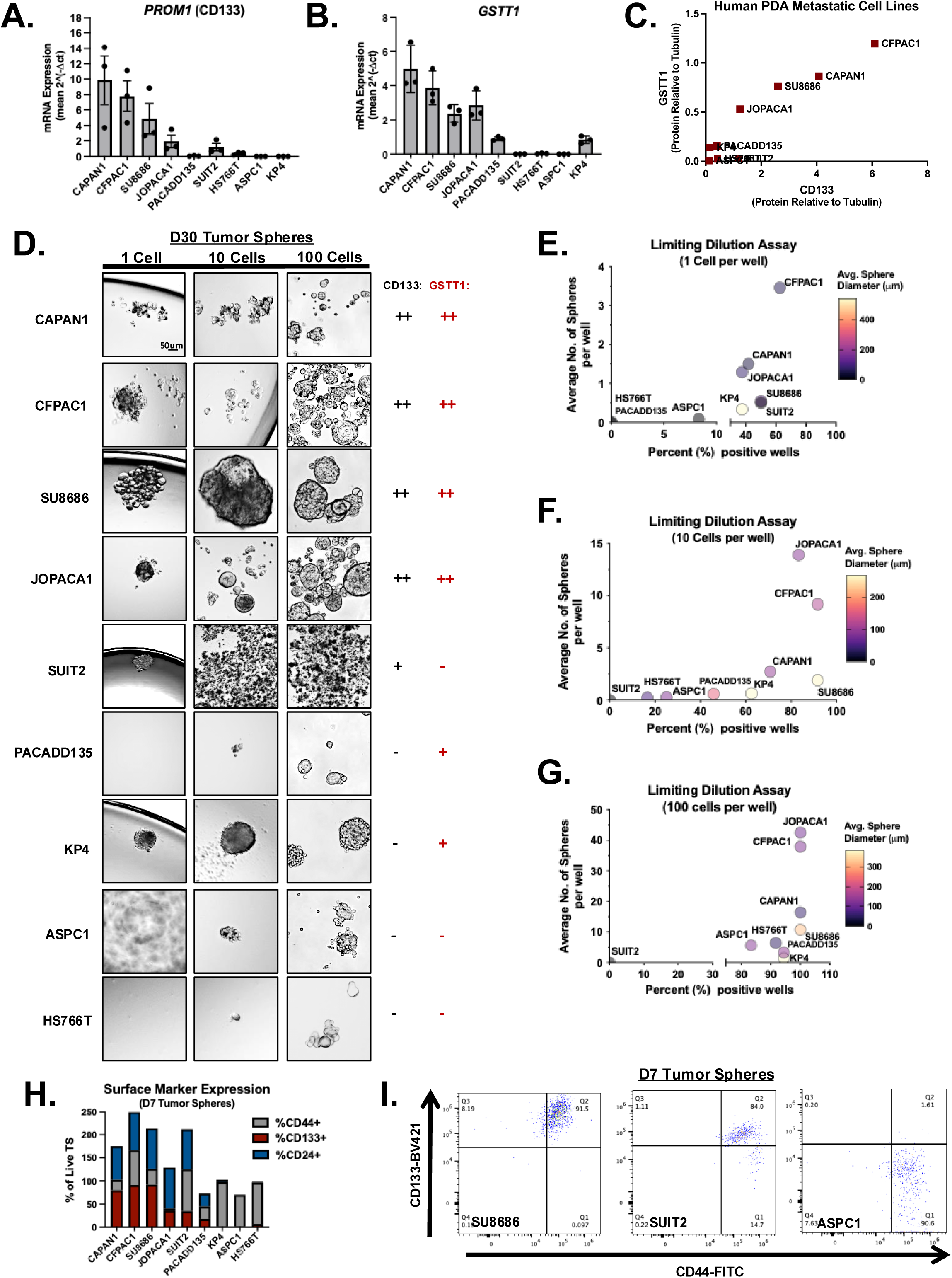

**Supp Fig 4.**
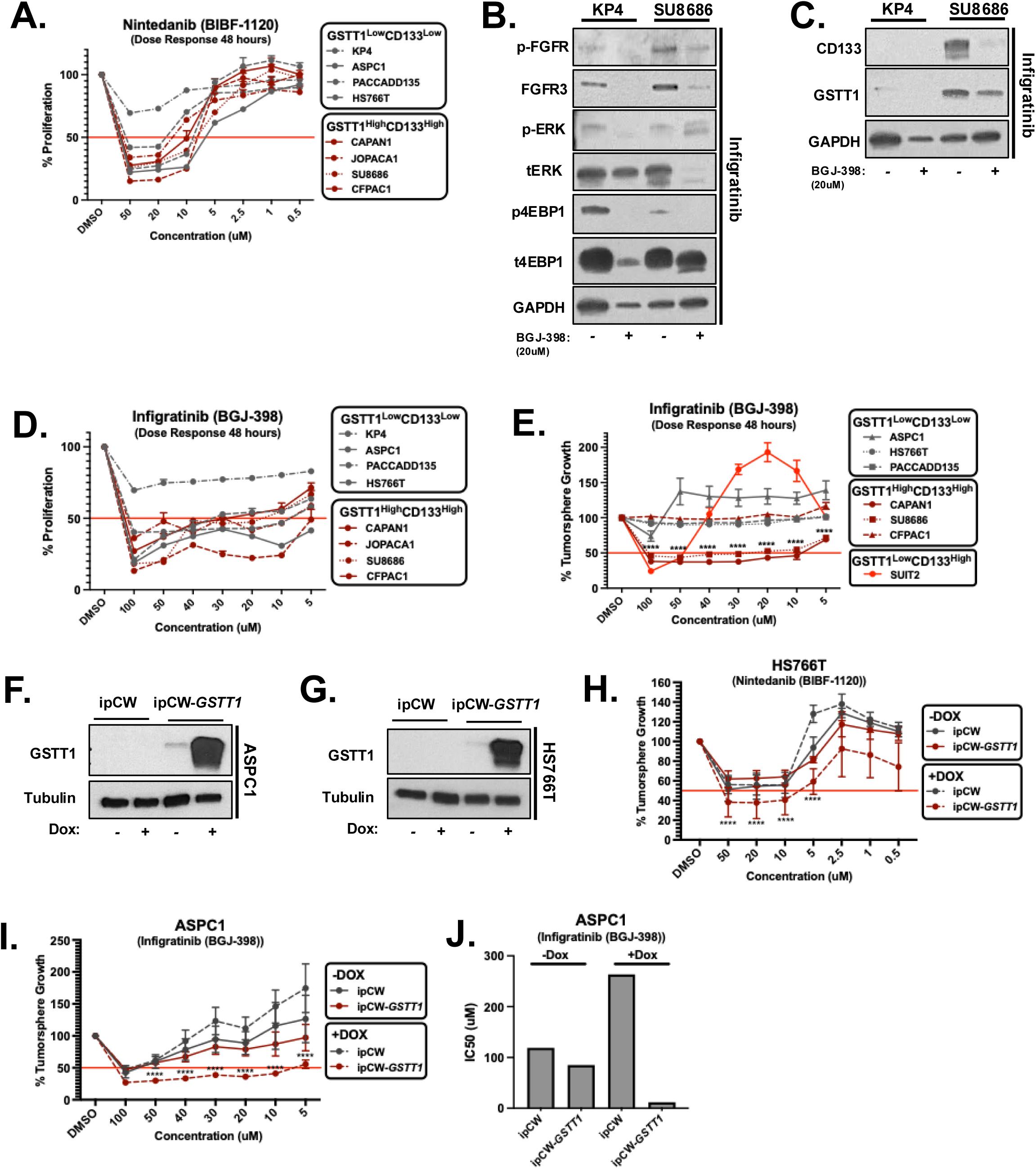

**Supp Fig 5.**
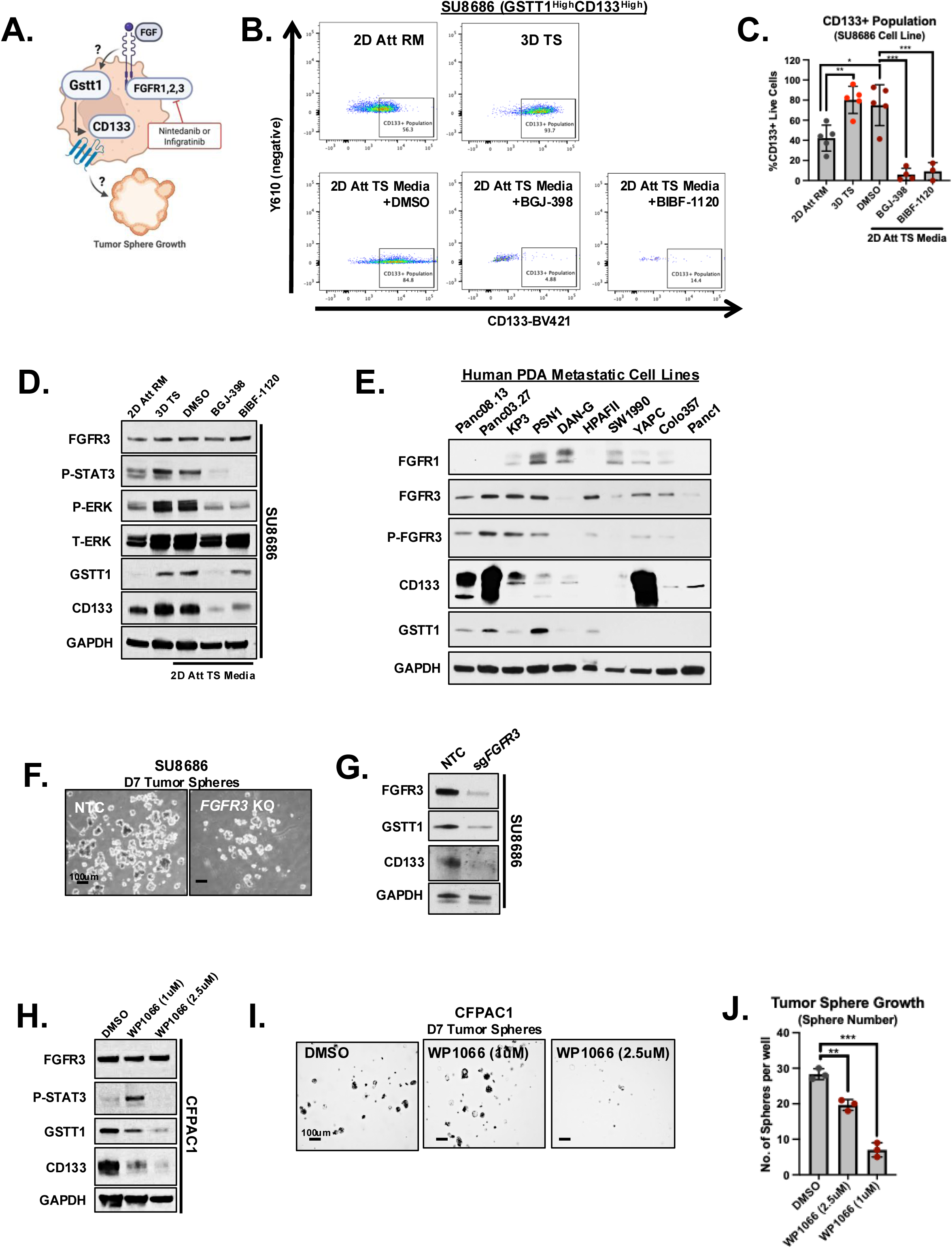

**Supp Fig 6.**
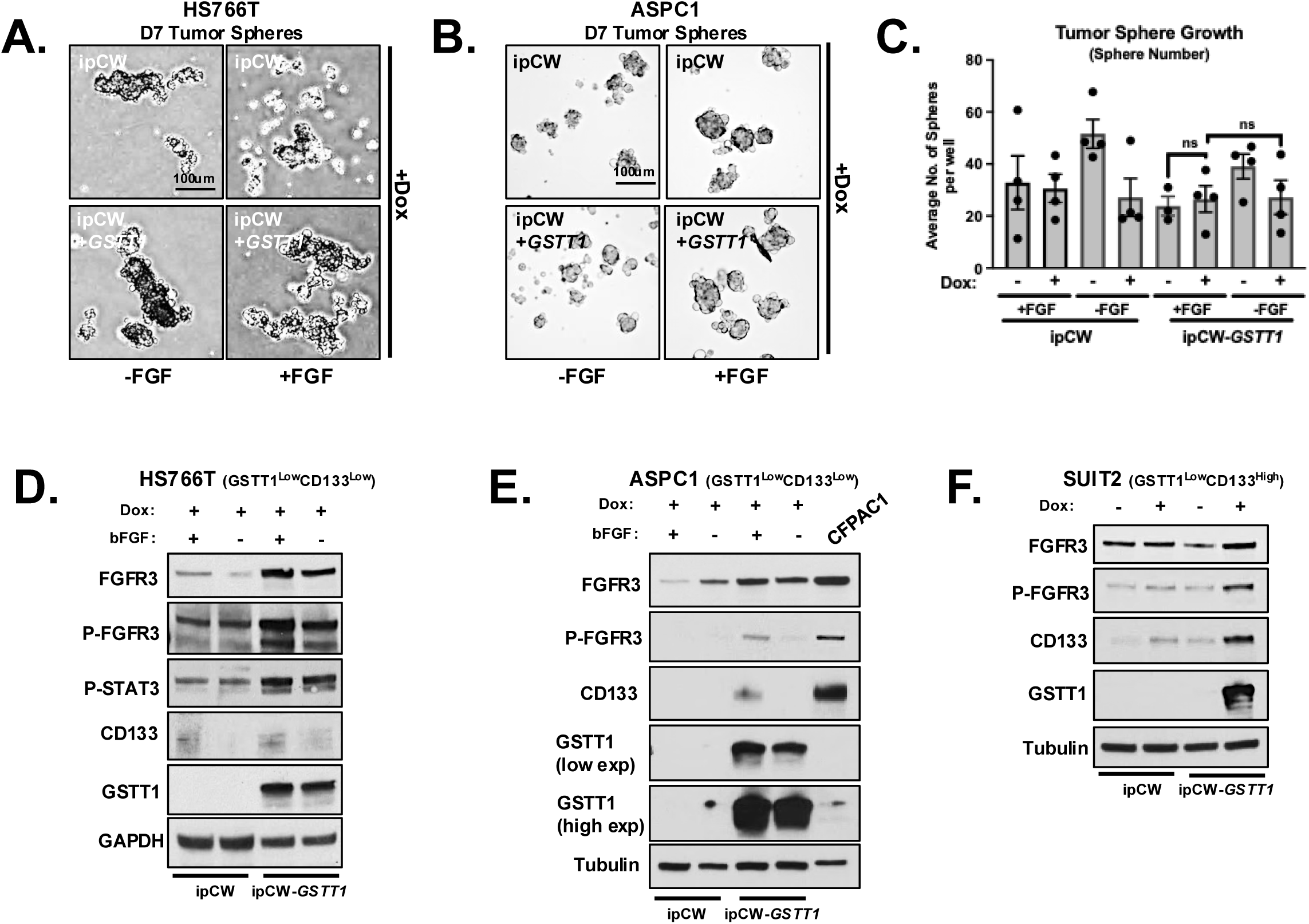

**Supp Fig 7.**
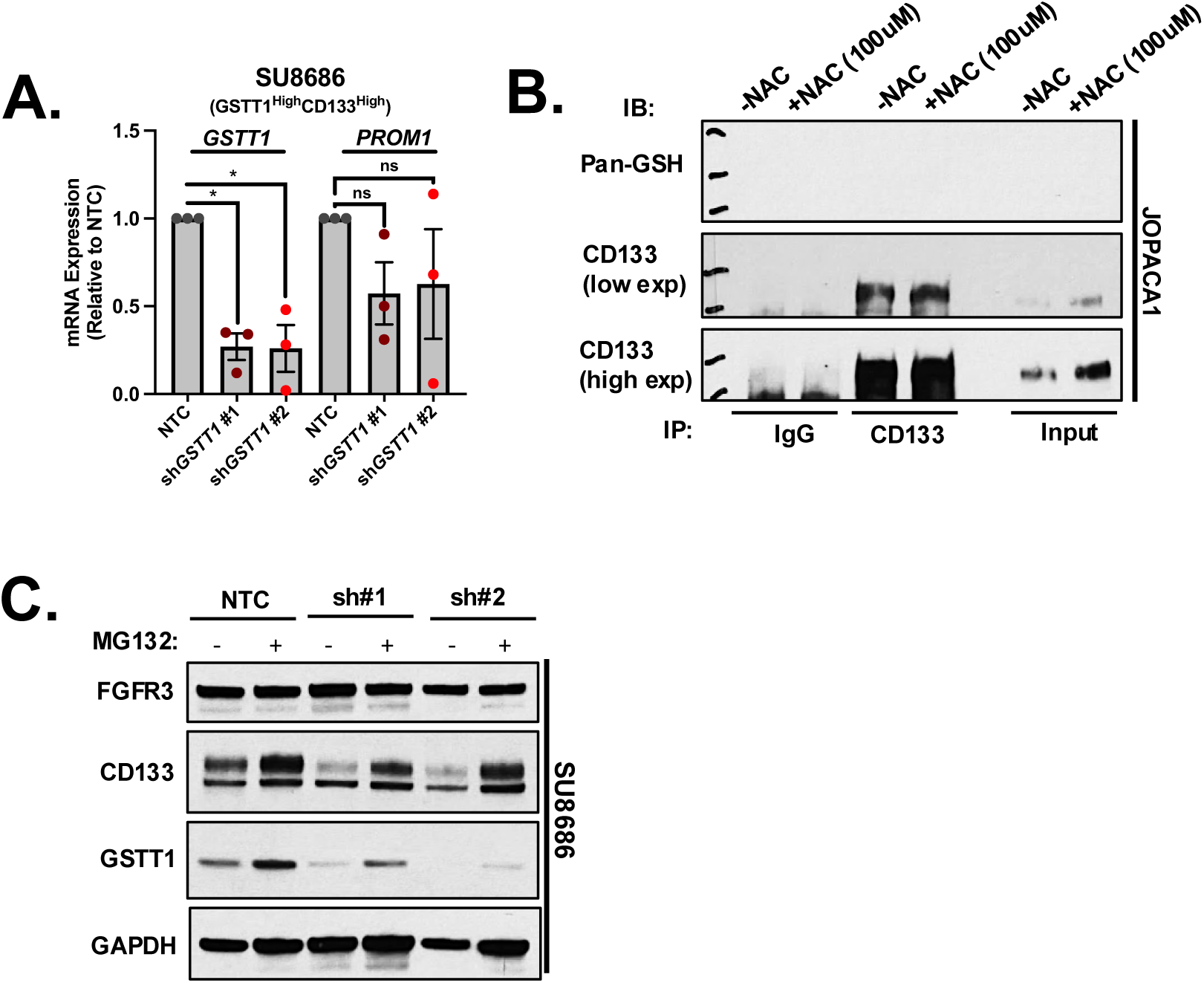

