## Supplementary Figure Legends for "GSTT1 mediates stemness and FGFR inhibitor sensitivity in pancreatic cancer through regulation of CD133 (*PROM1*)"

**Supplemental Figure 1. Gstt1^High^ Metastatic Tumor Spheres Retain mCherry Expression and are Enriched in Pathways Involved in GST Metabolism.** (A) Flow cytometry analysis of mCherry levels in Day 10 tumor sphere populations from Fig. 1. (B) Quantification of mCherry fluorescence in mCherry^High^ compared to mCherry^Low^ tumor spheres. Data represent N=2 independent experiments, with N=3 replicates each and are represented as mean s.d. (C) Ingenuity pathway analysis (IPA) of all differentially expressed genes identified in mCherry^High^ and mCherry^Low^ tumor spheres. Results are shown as -log *P*-value.

**Supplemental Figure 2. Identification of Human Pancreatic Cancer Cell Lines and Gstt1^High^ Tumor Sphere-Associated Survival Signatures.** (A) Schematic depicting the identification of relevant candidate genes in human pancreatic cancer from differentially expressed genes identified in mCherry tumor sphere populations. Created with Biorender.com. mCherry^High^ and mCherry^Low^ tumor cell populations were sorted and grown as tumor spheres for 10 days. Tumor spheres were then subjected to bulk RNA-Seq. N=106 upregulated genes identified in mCherry^High^ tumor spheres were analyzed for expression across human pancreatic cancer cell lines using the Cancer Cell Line Encyclopedia (CCLE). Out of n=106 genes, n=63 were significantly expressed in human pancreatic metastatic cell lines. n=63 genes were subsequently analyzed for differential effects on overall survival (OS) and relapse-free survival (RFS) in pancreatic cancer patients (KMplot). (B) n=37 genes were found to have differential prognosis on OS, A Mantel-Haenszel log-rank test was used to determine significance, **P*<0.05.

**Supplemental Figure 3. *PROM1* and *GSTT1* expression associate with tumor sphere initiation and growth in metastatic pancreatic cancer cells.** (A,B) qRT–PCR analysis of ***PROM1*** (A) and ***GSTT1*** (B) mRNA expression in a panel of human metastatic pancreatic cancer cell lines. Data are presented as mean ± s.e.m. from **N=3 independent experiments.** (C) Quantification of GSTT1 and CD133 protein levels from the western blots shown in Fig. 2B, normalized to tubulin loading control. (D) Representative bright-field images from tumor sphere limiting dilution assays performed in all nine cell lines (1, 10, and 100 cells per well), imaged 30 days after seeding. Scale bar for all, 50 µm. (E–G) Quantification of limiting dilution tumor sphere assays at **1 cell per well** (E), **10 cells per well** (F), and **100 cells per well** (G). The Y-axis represents the average number of tumor spheres per well, and the X-axis represents the percentage of positive wells containing tumor spheres. Bubble plots depict average tumor sphere diameter (µm), represented by color intensity. A minimum of **20-24 wells** were analyzed per condition. (H) Flow cytometry analysis of CD133, CD44, and CD24 surface marker expression across cell lines, shown as the percentage of the total Day 7 tumor sphere population. Data represent **N=2 independent experiments**. (I) Representative flow cytometry plots showing CD133 and CD44 surface expression in SU8686, SUIT2, and ASPC1 tumor spheres.

**Supplemental Figure 4. CD133^High^GSTT1^High^ Metastatic Tumor Spheres are Sensitive to FGFR Inhibitors.** (A) MTT dose response of 2D attachment metastatic pancreatic cancer cell lines treated with escalating doses of BIBF-1120 (Nintedanib) for 48 hours. Results depicted as % of DMSO control for each condition. Data represent **N=2 independent experiments** with N=3 replicates each, error bars indicate standard deviation (s.d.). (B, C) GSTT1^Low^CD133^Low^ (KP4) and GSTT1^High^CD133^High^ (SU8686) cell lines were treated with either DMSO or 20 µM BGJ-398 (Infigratinib) for 48 hours. Cell lysates were subjected to western blotting using the indicated antibodies. (D) MTT dose response of 2D attachment metastatic pancreatic cancer cell lines treated with escalating doses of BGJ-398 (Infigratinib) for 48 hours. Results depicted as % of DMSO control for each condition. Data represent **N=2 independent experiments** with N=3 replicates each, error bars indicate standard deviation (s.d.). (E) Tumor sphere growth of each cell line in response to BGJ-398, stratified by GSTT1 and CD133 levels. Results depicted as % of DMSO control for each cell line. Data represent N=3 independent experiments with N=3 replicates each, error bars indicate standard deviation (s.d.). Two-sided *t-test* with Welch’s correction was used to determine statistical significance between GSTT1^High^CD133^High^ cell lines compared to GSTT1^Low^CD133^Low^ cell lines for each dose (*****P*<0.0001). (F) ASPC1 and (G) HS766T were induced with doxycycline to express ipCW control or ipCW-*GSTT1* construct. Lysates were subjected to western blotting using indicated antibodies. Day 5 HS766T (H) and ASPC1 (I) tumor spheres induced with doxycycline to express ipCW control or ipCW-*GSTT1* construct were treated with escalating doses of BIBF-1120 or BGJ-398, respectively. Results depicted as % of DMSO control for each condition. Data represent N=3 independent experiments with N=3 replicates each, error bars indicate mean ± s.e.m. Two-sided *t-test* with Welch’s correction was used to determine statistical significance between ipCW-*GSTT1* +Dox condition was compared to all other conditions for each dose (***P*<0.01). (J) IC50 values for BGJ-398 (µM) in ASPC1 cell line for each condition were calculated from an average of N=3 independent experiments.

**Supplemental Figure 5. GSTT1 cooperates with FGF–FGFR signaling to enrich CD133 expression and tumor sphere growth in metastatic pancreatic cancer cells.**
(A) Schematic illustrating a proposed model in which FGF–FGFR signaling regulates GSTT1 and CD133 expression to promote tumor sphere growth in a subset of metastatic pancreatic cancer cells. Created with BioRender. (B) Representative flow cytometry plots showing CD133 surface expression in SU8686 cells cultured under the following conditions: 2D attachment, 3D tumor spheres, 2D attachment in tumor sphere (TS) media, 2D attachment TS media + BGJ-398 (20 µM), and 2D attachment TS media + BIBF-1120 (10 µM). (C) Quantification of the percentage of CD133⁺ cells among live cells from the experiment shown in (B). Data represent N=5 independent experiments, error bars indicate standard deviation (s.d.). A two-sided t-test was used to determine statistical significance between indicated comparisons (**P* < 0.05, ***P* < 0.01, ****P* < 0.001). (D) Western blot analysis of indicated proteins in SU8686 cells cultured under the conditions described in (B). (E) Western blot of CD133, GSTT1, FGFR3, and phospho-FGFR3 (Y724) in a panel of human pancreatic cancer cell lines (N=10). (F) Representative bright-field images of tumor spheres formed by SU8686 cells expressing non-targeting (NT) control or *FGFR3* knockout (KO). (G) Western blot analysis of FGFR3, CD133, and GSTT1 protein levels in SU8686 cells expressing NT control or *FGFR3* KO. (H) Western blot analysis of CFPAC1 cells treated with the STAT3 inhibitor WP1066 (1 or 2.5 µM) or DMSO control. (I) Representative bright-field images of CFPAC1 tumor spheres following treatment with DMSO, 1 µM WP1066, or 2.5 µM WP1066. (J) ImageJ quantification of the average tumor sphere number per well from (I). Data represent **N = 3 independent experiments** with **N = 3 technical replicates** per condition, error bars indicate standard deviation (s.d.). Statistical significance was determined using a two-sided t-test with Welch’s correction (***P* < 0.01, ****P* < 0.001).

**Supplemental Figure 6. *GSTT1* and FGF cooperate to induce low levels of CD133 protein but are insufficient to promote tumor sphere growth in GSTT1^Low^CD133^Low^ metastatic pancreatic cancer cells.** HS766T and ASPC1 (GSTT1^Low^CD133^Low^) metastatic pancreatic cancer cells were cultured under 2D attachment conditions and treated with either vehicle control (dH₂O) or basic FGF (bFGF; 20 ng/mL) for 7 days, with concurrent doxycycline induction of either ipCW control or ipCW-*GSTT1* constructs. (A,B) Representative bright-field images of Day 7 tumor spheres formed by ASPC1 and HS766T cells under the indicated conditions. Scale bar, 100 µm. (C) ImageJ quantification of tumor sphere number per well on Day 7 in ASPC1 cells. Data are presented as mean ± s.e.m. from **N = 4 independent experiments**, each with **n = 3 technical replicates**. Statistical significance was assessed using a two-sided t-test with Welch’s correction (ns, not significant). HS766T (D), ASPC1 (E), and SUIT2 (F) whole-cell lysates were analyzed by western blotting for the indicated proteins. ASPC1 protein levels were compared with endogenous expression in CFPAC1 (GSTT1^High^CD133^High^) cells.

**Supplemental Figure 7. GSTT1 regulates CD133 protein stability without altering *PROM1* transcription.** (A) qRT–PCR analysis of ***GSTT1*** and ***PROM1*** mRNA expression in SU8686 (GSTT1^High^CD133^High^) cells expressing a non-targeting (NT) control vector or two independent ***GSTT1*** shRNAs. Data are presented as mean ± s.e.m. from **N=3 independent experiments**. Statistical significance was assessed using a two-sided t-test with Welch’s correction (**P* < 0.05) and (ns). (B) Immunoprecipitation (IP) of IgG control or CD133 from JOPACA1 (GSTT1^High^CD133^High^) cells treated with glutathione via **N-acetylcysteine (NAC; 100 µM)**. Immunoprecipitates were immunoblotted for glutathione (GSH) and CD133. (C) SU8686 cells expressing NT control or two independent ***GSTT1*** shRNAs were treated in the presence or absence of the proteasome inhibitor **MG132 (5 µM),** followed by western blot analysis for the indicated proteins.
